# A previously unappreciated class of metal-dependent bile salt hydrolases from the human gut microbiome

**DOI:** 10.64898/2026.04.05.716592

**Authors:** Zheng Cui, Christopher J. Meng, Stephania M. Irwin, Hannah E. Augustijn, Panagiotis Panagopoulos Papageorgiou, Anh T. P. Nguyen, Ruocheng Yu, Miguel A. Aguilar Ramos, Heather J. Kulik, Emily P. Balskus

## Abstract

Bile salt hydrolases (BSHs) are gut microbial enzymes that catalyze the deconjugation of glycine-or taurine-conjugated bile acids (BAs), a key step in shaping the BA pool in the human gastrointestinal tract and modulating host-gut microbiome interactions.^1–3^ All known BSHs are members of the N-terminal nucleophile (Ntn) hydrolase superfamily and share a conserved architecture and mechanism involving a nucleophilic active site cysteine.^4,5^ This knowledge has guided predictions and study of BSH activity in the gut microbiome^6,7^ as well as the development of BSH inhibitors^8^. Here, we report the discovery and characterization of a previously unknown BSH from the human gut bacterium *Bilophila wadsworthia* that belongs to the metal-dependent amidohydrolase superfamily and exhibits robust and specific activity toward taurine-conjugated bile salts. We show this secreted enzyme, metalloBSH, utilizes a metallocofactor for BA deconjugation, a mechanism distinct from that of canonical Ntn-type BSHs. MetalloBSHs are conserved in *B. wadsworthia* and present in many other *Desulfovibrionaceae* found in vertebrate gut microbiomes. Analysis of multi-omic datasets indicates metalloBSHs are expressed in vivo and correlate with BA metabolism. Overall, our findings reshape our understanding of BSH activity in the gut microbiome and highlight the promise of activity guided discovery in revealing previously overlooked gut microbial enzymes.

## Introduction

The primary BAs cholic acid (CA) and chenodeoxycholic acid (CDCA) are synthesized in the human liver from cholesterol and conjugated with glycine or taurine to form conjugated primary BAs or bile salts that are excreted into the small intestine.^9^ A fraction of conjugated BAs is not reabsorbed and undergoes extensive chemical modification by gut microbes to form a wide variety of secondary BAs.^10–12^ BAs play multiple important roles in human physiology,^13^ aiding in digestion and nutrient absorption,^14^ shaping the composition of the gut microbiome,^15^ and functioning as signaling molecules to regulate numerous host processes.^16–18^

Gut bacterial BSH enzymes catalyze the first step in conjugated primary BA metabolism by hydrolyzing the amide bond of conjugated BAs, releasing glycine or taurine.^11^ Currently, all known BSHs belong to the N-terminal nucleophile (Ntn) hydrolase superfamily, which uses a conserved N-terminal nucleophilic cysteine residue to catalyze amide bond hydrolysis (**Fig. 1a**).^2^ These enzymes were also recently found to catalyze amide bond formation between primary BAs and additional amino acids.^19,20^ Because of the importance of BA metabolism to the gut microbiome and human health, BSHs have been the focus of extensive study. For example, profiling of Human Microbiome Project (HMP) cohort metagenomes has revealed more than 590 *bsh* genes assigned to 117 gut bacterial genera from 12 phyla.^7,21^ BSH activity has been connected to both positive and negative health outcomes.^3,6,22^ Finally, small molecule inhibitors of BSHs have been developed and used as tools.^8^ Notably, our current understanding of its distribution, function, health relevance, and inhibition assumes this family of enzymes is solely responsible for BSH activity.

**Fig. 1.**
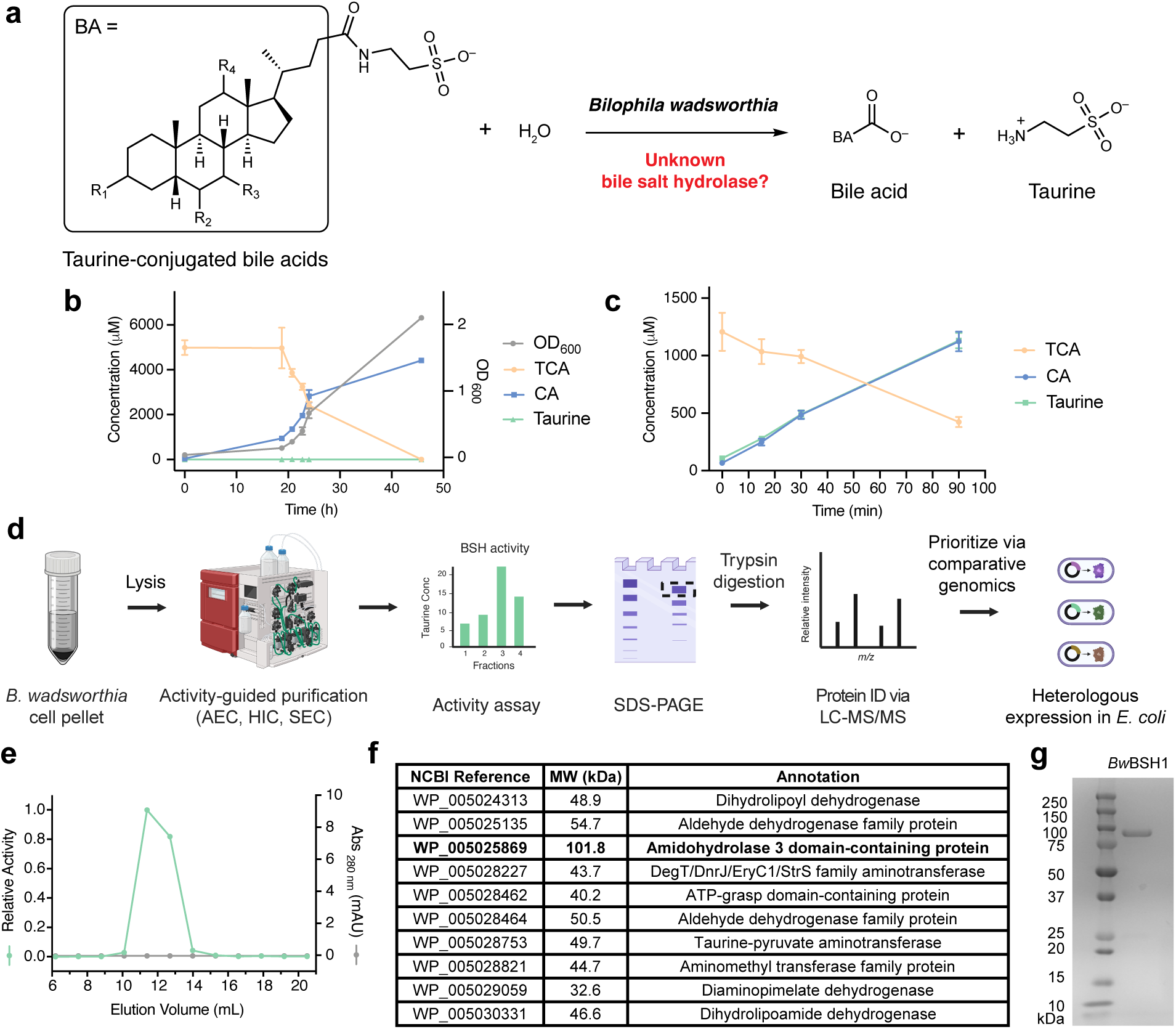
Activity guided purification reveals a bile salt hydrolase from *Bilophila wadsworthia* (*Bw*BSH). a, BSH catalyzes the hydrolysis of bile salts, including taurine-conjugated bile acids. b, Measurement (OD_600_) of *B. wadsworthia* 3_1_6 growth in anaerobe basal broth (ABB) supplemented with 5 mM TCA paired with LC-MS/MS analysis of levels of taurine, TCA, and CA shows hydrolysis of TCA correlated with growth. c, *B. wadsworthia* cell suspension deconjugates TCA. For b and c, data points represent the mean of N = 3 biological replicates. Error bars indicate standard deviation. d, Activity-guided *Bw*BSH discovery workflow. e, Relative TCA hydrolysis activity and UV absorption at 280 nm of the indicated fraction following anion-exchange chromatography (AEC), hydrophobic interaction chromatography (HIC) and size-exclusion chromatography (SEC). f, Candidate BSHs chosen for heterologous expression with active BSH (*Bw*BSH1) bolded. g, SDS-PAGE gel image of *Bw*BSH1 purified from recombinant *E. coli* BL21(DE3) cultures.

*Bilophila wadsworthia* is a sulfite-reducing intestinal bacterium that generates hydrogen sulfide from metabolism of taurine and isethionate.^23,24^ It has been implicated in diseases ranging from inflammatory bowel disease (IBD) to colorectal cancer (CRC).^25–28^ *B. wadsworthia* has long been connected to BAs due to its ability to grow in the presence of high concentrations of conjugated BAs and its use of taurine as an energy source. Correlations between this gut bacterial species and BAs have also been observed in vivo. Prior work demonstrated that supplementation of taurocholic acid (TCA) in a low-fat diet promotes the expansion of *B. wadsworthia* and development of colitis in *Il10^-/-^* mice.^26^ Metabolomic and microbiome sequencing data from clinical studies revealed that the relative abundance of *B. wadsworthia* correlates with altered host gut BA pools, specifically showing an inverse relationship with taurine-conjugated BAs^29^ and a positive association with deconjugated primary BAs^30^. Finally, a human gut isolate that shares 99.6% 16S rDNA nucleotide sequence similarity with *B. wadsworthia* was reported to deconjugate TCA both in vitro and in gnotobiotic mice.^31^ However, the genomes of *B. wadsworthia* strains lack genes encoding recognizable BSH homologs^29^, leaving this potential enzymatic activity unexplained.

## Results

### Bilophila wasworthia hydrolyzes TCA

To explore the ability of *B. wadsworthia* to hydrolyze taurine-conjugated BAs, we initially used a growth-based assay. *B. wadsworthia* 3_1_6 was cultured anaerobically in supplemented anaerobe basal broth (ABB). We confirmed that ABB supplemented with taurine supported growth. In contrast, neither ABB alone nor ABB supplemented with glycocholic acid (GCA) permitted growth. Notably, supplementation of ABB with either TCA or taurodeoxycholic acid (TDCA) also supported the growth of *B. wadsworthia* (**Supplementary Fig. 1**), and this growth correlated with TCA consumption and cholic acid production as measured by Ultra High-Performance Liquid Chromatography Tandem Mass Spectrometry (UPLC-MS/MS) analysis. This result indicated that *B. wadsworthia* can deconjugate TCA and release taurine (**Fig. 1b**).

To assess whether this TCA deconjugation activity arises from enzymes produced by *B. wadsworthia*, we grew *B. wadsworthia* in ABB supplemented with taurine, pelleted the bacteria, washed the cell pellet with phosphate-buffered saline (PBS) and resuspended the cells in PBS containing TCA, TDCA, or GCA under aerobic conditions, which prevents growth. We observed the deconjugation of TCA and TDCA in the resting cell suspensions (**Fig. 1c**, and **Supplementary Fig. 2**). No activity was detected in assays containing heat-treated cells or a PBS control, consistent with enzymatic catalysis. The lack of metabolism of GCA suggested that putative BSH from *B. wadsworthia* specifically hydrolyzes taurine- but not glycine-conjugated BAs. This assay also suggested that, like canonical BSHs, the putative BSH in *B. wadsworthia* is insensitive to oxygen.

As highlighted above, *B. wadsworthia* genomes lack genes encoding canonical BSHs. We initially attempted to identify *B. wadsworthia* species that are incapable of BA deconjugation to enable BSH identification via comparative genomics. However, all three *B. wadsworthia* strains commercially available at the time of this experiment (ATCC 49260, *B. wadsworthia* 3_1_6 and *Bilophila* sp. 4_1_30) grew in medium containing TCA, indicating that they all possess BSH activity (**Supplementary Fig. 3**). Next, we assessed the inducibility of BSH activity in *B. wadsworthia* to gauge the feasibility of using comparative transcriptomics or proteomics approaches for enzyme identification. We grew cultures in ABB containing sulfite as a terminal electron acceptor and supplemented growing cultures with either TCA or PBS. If BSH expression was inducible, cell suspensions from TCA-supplemented cultures should hydrolyze TCA more rapidly than cell suspensions from cultures supplemented with PBS. However, cell suspensions from both cultures depleted TCA and produced taurine and CA at similar rates (**Supplementary Fig. 4**), indicating that the BSH in *B. wadsworthia* is likely constitutively expressed.

#### Activity-guided protein purification reveals a distinct BSH from *B. wadsworthia*

Given these results, we employed activity-guided native protein purification to identify the BSH from *B. wadsworthia* 3_1_6 (**Fig. 1d**), fractionating culture lysates and measuring TCA hydrolysis activity through multiple purification steps before a final purification using size exclusion chromatography (**Fig. 1e**). MS analysis of tryptic peptides from two active fractions revealed peptides that mapped to 75 proteins in the *B. wadsworthia* 3_1_6 proteome (**Supplementary Fig. 5**, **Supplementary Table 1**). Proteins detected in both active fractions were prioritized. To further narrow candidates for experimental evaluation, we employed comparative genomics with *E. coli* K-12 MG1655 (GCA_904425475.1). Because *E. coli* lacks TCA deconjugation activity, *B. wadsworthia* hit proteins with > 36% amino acid (aa) sequence identity to *E. coli* proteins were deprioritized. This left 10 candidate BSHs (**Fig. 1f**) which we heterologously expressed individually in *E. coli* as N-hexahistidine (His_6_)-tagged fusions, purified, and tested for BSH activity toward TCA. We observed high BSH activity only with protein WP_005025869 (UniProt ID: E5Y4F1), identifying it as the likely BSH from *B. wadsworthia* 3_1_6 (*Bw*BSH1) (**Fig. 1g**).

*Bw*BSH1 is a 101 kDa protein annotated as a metal-dependent amidohydrolase and includes a primary Amidohydro_3 domain (Pfam: PF07969, *E* = 2.4 × 10^-63^)^32^. Members of this enzyme superfamily, including urease^33^, phosphotriesterase^34^, dihydroorotase^35^, hydantoinase^36^, and carboxypeptidase^37^, share a conserved TIM barrel structural fold and a mononuclear or dinuclear metal center (typically Zn^2+^, Mn^2+^, Fe^2+^ or Ni^2+^). The metal center is ligated by structurally conserved amino acids, which typically include histidine, aspartate, and/or glutamate residues^38^. The metallocofactor coordinates and activates a water molecule for nucleophilic attack on a bound electrophile. Though characterized members of this superfamily catalyze a variety of hydrolytic transformations, to date, no amidohydrolases have been reported to possess BSH activity. Sequence comparison of *Bw*BSH1 and a canonical Ntn-BSH from *Bacteroides thetaiotaomicron* (BT_2086, NCBI Reference: WP_008761025) revealed no primary aa sequence similarity (9.6% pairwise identity) and the absence of the catalytic N-terminal Cys. Pairwise structure alignment of an AlphaFold3^39^-predicted structure of *Bw*BSH1 with the crystal structure of Ntn-BSH from *B. thetaiotaomicron* (PDB: 6UFY)^40^ showed minimal structural similarity with a TM (Template Modeling) score^41^ of 0.0917. The lack of protein sequence and structural similarity to Ntn-BSHs explains why *Bw*BSH was not readily identified using computational approaches.

To experimentally test the potential involvement of a catalytic Cys in catalysis by *Bw*BSH1, we evaluated the activity of the pan Ntn-BSH inhibitor BSH-IN-1^40^ toward the purified enzyme. BSH-IN-1 bears an α-fluoromethyl ketone warhead that covalently modifies the catalytic cysteine residue of these enzymes. We preincubated 8 nM *Bw*BSH1 with varying concentrations of BSH-IN-1 (100 nM–1000 µM) for 30 min and then assayed TCA hydrolysis. Notably, unlike its potent inhibition of canonical Ntn-BSHs (IC_50_ = 427 nM for BT_2086), BSH-IN-1 slightly enhanced *Bw*BSH1 activity in a concentration-dependent manner (**Fig. 2a**). This observation suggests that *Bw*BSH1 employs a catalytic mechanism distinct from canonical Ntn-BSHs.

**Fig. 2.**
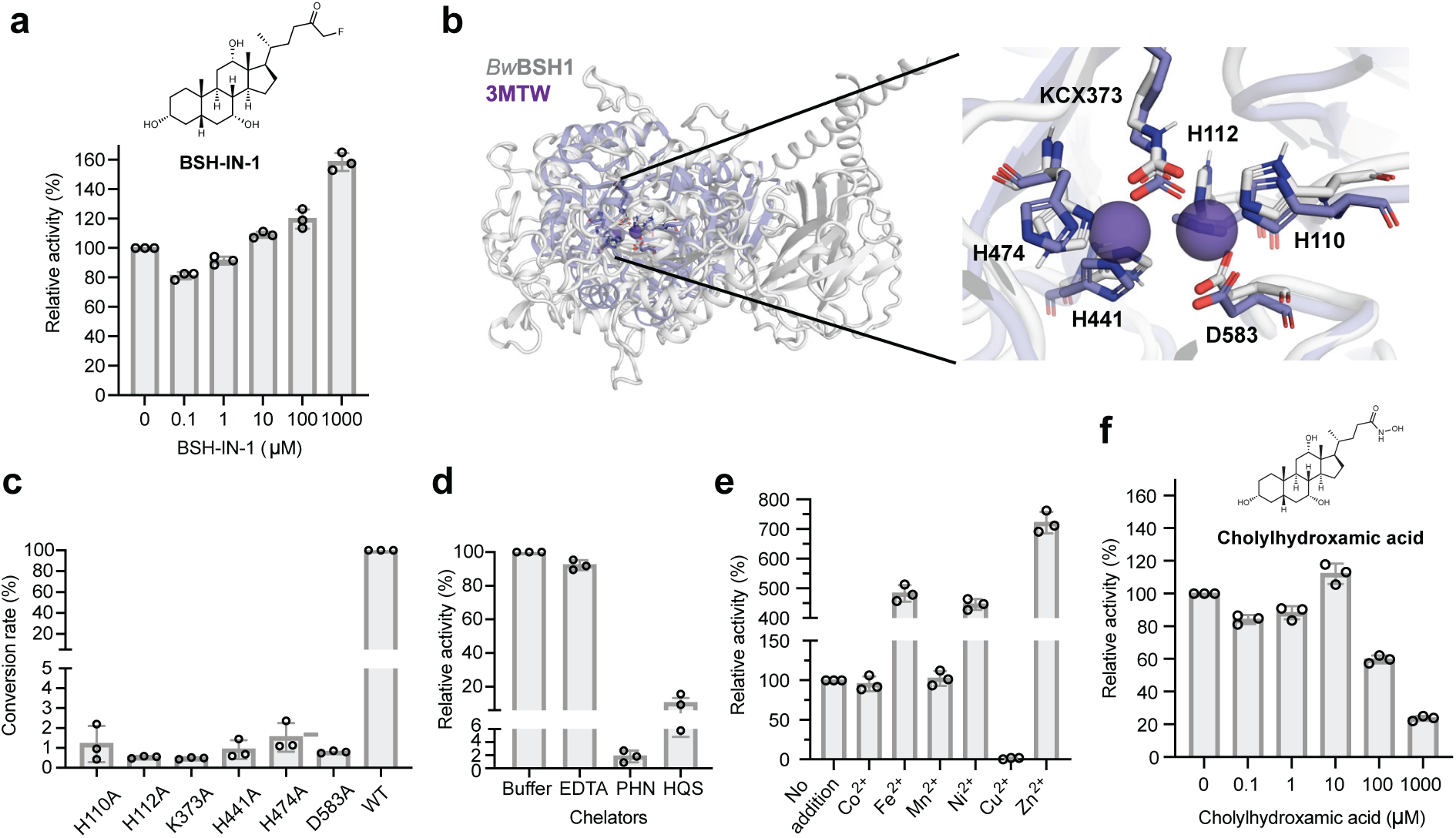
***Bw*BSH is a metal-dependent enzyme**. a, Evaluation of pan-BSH inhibitors on *Bw*BSH1 activity using a UPLC-MS/MS end-point assay. b, Superimposition of the AlphaFold3-predicted *Bw*BSH1 structure (grey) onto the crystal structure of a zinc-dependent dipeptidase from *Caulobacter crescentus* (PDB: 3MTW; purple). The two bound Zn^2+^ ions are retained from 3MTW, with the corresponding conserved metal-coordinating residues (H110, H112, KCX373, H441, H474, and D583) labeled in the *Bw*BSH1 model. c, Impact of site-directed mutagenesis of conserved, putative metal-binding residues on *Bw*BSH1 activity. Conversion rate (%) is expressed as the percentage of taurine product generated by each mutant compared to wild-type *Bw*BSH1 following an overnight incubation. d, *Bw*BSH1 hydrolysis activity at 37 °C following a 16 h pre-incubation at 4 °C with buffer (control) or 5 mM of the metal chelators EDTA, 1,10-phenanthroline (PHN), or 8-hydroxyquinoline-5-sulfonic acid (HQS). e, Effect of metal ion addition on the activity of metal-depleted *Bw*BSH1 purified from *E. coli* grown in M9 minimal medium. f, Dose-dependent inhibition of *Bw*BSH1 by cholylhydroxamic acid, evaluated via an end-point activity assay. For panels a, d, e, and f, relative activity (%) was determined by normalizing the concentration of taurine product generated in enzyme-containing samples to that in the vehicle control under initial rate conditions. Assays were carried out using 5 nM *Bw*BSH1 and 1 mM TCA for 15 min to capture the relative activity under initial rate conditions. For all panels, bars represent mean values of three biological replicates (N = 3); error bars indicate standard deviation.

### *B. wadsworthia* BSHs are metalloenzymes

Given its annotation as a metal-dependent amidohydrolase, we next investigated the metal dependency of *Bw*BSH. To assess the presence of key metal-binding residues in *Bw*BSH, we analyzed its sequence and predicted structure. Aligning the sequences of *Bw*BSHs and characterized metal-dependent amidohydrolases using the Conserved Domains Database (CDD)^42^ revealed five conserved metal-coordinating residues and a conserved lysine residue that is often post-translationally carboxylated via spontaneous reaction with CO_2_ (**Extended Data Fig. 1**). Additionally, a Foldseek^43^ search against the PDB100 database, using the AlphaFold3-predicted apo structure of *Bw*BSH1 revealed that many of its structural homologs contain zinc ions (**Supplementary Table 2**). Structural alignment with its closest structural homologs, an uncharacterized metal-dependent hydrolase from *Pyrococcus furiosus* (PDB: 3ICJ) and an L-arginine carboxypeptidase from *Caulobacter crescentus* (PDB: 3MTW)^44^, further supported that these conserved active-site amino acids form a dinuclear zinc-binding metallocofactor. Specifically, a more buried Zn^2+^ ion (M_⍺_) is predicted to coordinate with H110, H112 and D583, while a more solvent exposed Zn^2+^ ion (M_β_) is predicted to coordinate with H441 and H474. Based on comparisons with these structural homologs and other metal-dependent hydrolases^38^, we predict that *Bw*BSH binds two metal ions that are bridged by a hydroxide ligand and a post-translationally modified carboxylated lysine (KCX373) (**Fig. 2b**). To test the importance of these putative metal-binding residues to *Bw*BSH activity, we expressed, purified, and tested the corresponding single-point alanine mutants (**Supplementary Fig. 6**). All mutations (H110A, H112A, D583A, K373A, H441A, and H474A) abolished BSH activity (**Fig. 2c**), consistent with an essential role for these amino acids in coordinating a dinuclear metallocofactor.

To identify the metal(s) present in *Bw*BSH1, we analyzed the protein using inductively coupled plasma mass spectrometry (ICP-MS). For this analysis, N-terminally His_6_-tagged *Bw*BSH1 was expressed in *E. coli* cultured in Lennox LB medium and purified via nickel-affinity chromatography. This revealed zinc, iron, nickel, and copper, with zinc observed in the greatest abundance (0.59 atoms per protein monomer) (**Supplementary Table 3**). Treating *Bw*BSH1 with the small molecule metal chelator EDTA revealed minimal reduction in activity but treatment with the more hydrophobic chelators 8-hydroxyquinoline-5-sulfonic acid (HQS) and 1,10-phenanthroline (PHN) substantially reduced activity (**Fig. 2d**). This differential susceptibility is consistent with inhibition profiles observed for other amidohydrolase superfamily enzymes^45–47^. In these related enzymes, bulky, hydrophilic chelators like EDTA are often unable to access buried active-site metal centers, whereas smaller, hydrophobic chelators like PHN and HQS can successfully access the active site and remove essential metals. Together, these results provide functional evidence that *Bw*BSH requires a tightly bound metallocofactor for catalytic activity.

To identify the specific metal(s) involved in catalysis, we prepared metal-depleted *Bw*BSH1 by expressing the enzyme in *E. coli* grown in M9 minimal medium and examined the effect of supplementing individual metal ions (Co^2+^, Fe^2+^, Mn^2+^, Ni^2+^, Cu^2+^, and Zn^2+^) on enzyme activity. By ICP-MS analysis, the metal-depleted *Bw*BSH1 contained < 0.11 metal ions per monomer of protein except for Ni^2+^ (0.78 ions per monomer), which likely carried over from the nickel-affinity purification step (**Supplementary Table 3**). Such metal carryover and mixed-metal incorporation are frequently observed when heterologously expressing amidohydrolase superfamily members in *E. coli*.^38,48^ Adding metals to metal-depleted *Bw*BSH had variable effects: Cu^2+^ caused strong inhibition, while Zn^2+^ most strongly enhanced activity (**Fig. 2e**). The capacity to utilize multiple different metal ions for catalysis is a feature of the amidohydrolase superfamily, and many members can be functionally reconstituted with various divalent cations (e.g., Co^2+^, Mn^2+^, Cd^2+^, Fe^2+^, or Zn^2+^).^48,49^ We also evaluated metal supplementation with chelator-treated *Bw*BSH1, which contained < 0.15 metal ions per protein monomer for all examined metals (**Supplementary Table 3**). This approach yielded similar results, with Cu^2+^ inhibiting and Zn^2+^ enhancing activity (**Extended Data Fig. 2a**). Metal supplementation had little impact on the activity of *Bw*BSH1 purified from LB medium, though Cu^2+^ consistently demonstrated an inhibitory effect (**Extended Data Fig. 2b**). Comparing the two metal-depletion methods revealed that both preparations exhibited substantially lower activity than *Bw*BSH1 purified from LB medium, with the M9-expressed enzyme showing less activity than the chelator-treated enzyme (**Extended Data Fig. 2c**), further underscoring the necessity of a metallocofactor. Finally, we prepared Zn^2+^- and Cu^2+^-reconstituted *Bw*BSH1 by incubating the metal-depleted enzyme with ZnCl_2_ or CuCl_2_, the metal ions exhibiting the most pronounced activity differences, followed by overnight dialysis to remove unbound metal ions. ICP-MS analysis showed that Zn^2+^-reconstituted *Bw*BSH contained 4 equivalents of Zn per monomer of protein and Cu(II)-reconstituted *Bw*BSH contained approximately 6 equivalents of Cu (**Supplementary Table 3**). While these results are potentially consistent with *Bw*BSH containing a dinuclear Zn^2+^ metallocofactor, additional work is required to precisely define its structure.

Finally, we hypothesized that, in contrast to canonical Ntn-BSHs, use of a metallocofactor should render *Bw*BSH1 susceptible to inhibition by BA analogs bearing metal chelating functional groups. To test this hypothesis, we synthesized cholylhydroxamic acid (CHA)^50,51^ (**Fig. 2f**), which contains a hydroxamic acid capable of coordinating to a metallocofactor. We found that CHA inhibited *Bw*BSH1 at concentrations > 100 µM (**Fig. 2f**). Taken together, these data indicate that *Bw*BSH employs a metallocofactor to catalyze bile salt hydrolysis, further supporting its assignment as a metal-dependent BSH (metalloBSH).

### *B. wadsworthia* secretes multiple metalloBSHs

Further examination of the protein sequence of *Bw*BSH1 revealed additional features that distinguish this enzyme from canonical Ntn-BSHs. While Ntn-BSHs contain 220–420 aa and consist of a single protein domain, *Bw*BSH1 is 945 aa in length and, along with the metal-dependent hydrolase domain, is predicted to contain a high-confidence N-terminal signal peptide (SignalP 6.0^52^ likelihood = 0.9903) for Sec translocon-mediated export, as well as a C-terminal domain and a C-terminal disordered region (**Fig. 3a**). Examining the genomic context of *Bw*BSH1 in *B. wadsworthia* 3_1_6 revealed a second gene encoding a homolog of *Bw*BSH1 with 50.9% aa identity (*Bw*BSH2, WP_016360761, UniProt ID: E5Y4F0) located adjacent to genes encoding a predicted type II secretion system (T2SS).^53^ Analysis of the genomes of the two additional commercially available *B. wadsworthia* strains (*Bilophila* sp. 4_1_30, and *B. wadsworthia* ATCC 49260) revealed similar genetic loci, with *Bilophila* sp. 4_1_30 encoding two colocalized *Bw*BSHs and *B. wadsworthia* ATCC 49260 encoding three colocalized enzymes (**Fig. 3b**).

**Fig. 3.**
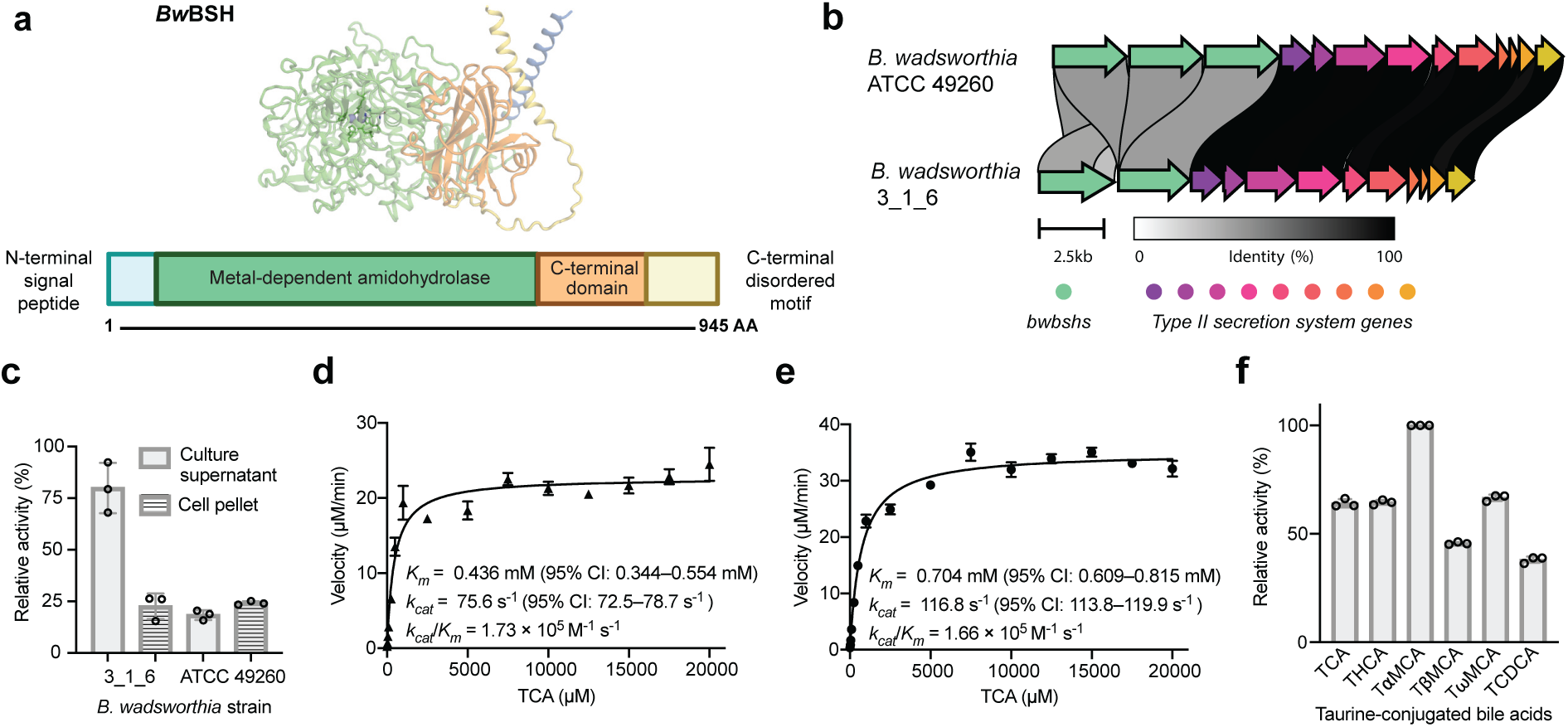
Secreted *Bw*BSHs possess high activity and specificity toward taurine-conjugated BAs. a, Domains of a *Bw*BSH. b, Genome neighborhoods of *bsh*s from *B. wadsworthia* 3_1_6 and ATCC 49260. c, TCA hydrolysis activity is detected in *B. wadsworthia* culture supernatants and cell pellets by incubating with 1 mM TCA at 37 °C for 15 min and measuring the initial hydrolysis activity. d, e, Single-substrate kinetic analyses of TCA hydrolysis with 5 nM recombinant *Bw*BSH1 (d) and 5 nM recombinant *Bw*BSH2 (e). f, Relative activity of *Bw*BSH1 towards 1 mM taurine-conjugated bile acids including TCA, taurohyocholic acid (THCA), tauro-⍺-muricholic acid (T⍺MCA), tauro-β-muricholic acid (TβMCA), tauro-ω-muricholic acid (TωMCA), and taurochenodeoxycholic acid (TCDCA). Data represent mean values of three biological replicates (N = 3); error bars indicate standard deviation.

The presence of a predicted signal peptide in *Bw*BSHs and the adjacent predicted secretion system led us to hypothesize that, unlike canonical Ntn-BSHs which are often cytoplasmic*, Bw*BSH enzymes are secreted. To test this, we measured BSH activity in both culture supernatants and cell lysates of *B. wadsworthia* 3_1_6 and *B. wadsworthia* ATCC 49260 (**Fig. 3c**). We detected higher activity in the supernatants compared to cell lysates of *B. wadsworthia* 3_1_6. We then purified secreted native *Bw*BSH from the medium of TCA-supplemented cultures using an activity guided approach (**Extended Data Fig. 3a**). MS analysis of tryptic peptides from the purified protein confirmed the presence of both *Bw*BSH1 and *Bw*BSH2 (**Extended Data Fig. 3b, c**, and **Supplementary Table 4**), indicating both enzymes are expressed and active. At equivalent enzyme concentrations, the activity of the natively purified *Bw*BSH enzymes was comparable to that of recombinant *Bw*BSH1 expressed in *E. coli* BL21(DE3) (**Extended Data Fig. 3d, Supplementary Fig. 7**). These findings support the hypothesis that *B. wadsworthia* secretes multiple BSHs to facilitate bile salt hydrolysis outside the cell, releasing taurine to support its growth (**Supplementary Fig. 8**).

To further assess the activities of *Bw*BSHs, we heterologously expressed *Bw*BSH1 and *Bw*BSH2 individually in *E. coli*, purified each enzyme, and determined their activities using TCA as a substrate in PBS buffer at pH 7.4 and 37 °C. Single-substrate kinetic analysis revealed typical Michaelis–Menten kinetic parameters for both enzymes (*Bw*BSH1 *k_cat_*/*K_m_* = 1.73×10^5^ M^-1^ s^-1^, *Bw*BSH2 *k_cat_*/*K_m_* = 1.66×10^5^ M^-1^ s^-1^) (**Fig. 3d, e**). The catalytic efficiencies fall at the upper end of the range reported for canonical Ntn-BSHs, which typically have *k_cat_/K_m_* values of 3×10^3^ – 8×10^5^ M^-1^ s^-1^ at pH 6 and 37 °C (**Supplementary Table 5**)^54–56^. In contrast to canonical Ntn-BSHs, *Bw*BSHs display a broader pH optimum spanning pH 4.0–5.4 and retain ∼14% of their maximal initial activity at pH 8.0 (**Supplementary Fig. 9**).

Examining the substrate specificity of *Bw*BSH1 revealed high catalytic efficiencies toward multiple taurine-conjugated BAs tested, with maximal activity observed for tauro-⍺-muricholic acid (**Fig. 3f**). In addition, *Bw*BSH1 hydrolyzed several other *N*-acyl taurines, albeit less efficiently (**Supplementary Fig. 10a**). Consistent with our initial culture-based assays, no activity was observed toward glycine-conjugated BA (**Supplementary Fig. 10b**) . Unlike canonical Ntn-BSHs, which exhibit acyltransferase activity and catalyze the formation of amine-conjugated bile acids, *Bw*BSH1 could not conjugate amino acids to cholic acid. Altogether, these experiments demonstrate that *Bw*BSHs have high activity and specificity for the deconjugation of taurine-conjugated BAs, features that are consistent with these metalloBSHs contributing to taurine availability for *B. wadsworthia* metabolism.

#### Delineation of the metalloBSH enzyme family

We next sought to uncover features that distinguish metalloBSHs from other metal-dependent amidohydrolases to understand their unique activity and assess their distribution in microbial genomes and metagenomes. To identify amino acids beyond the conserved metal-binding residues that may mediate substrate recognition and BSH activity, we performed docking studies using the AlphaFold3-predicted structure of Zn^2+^-containing *Bw*BSH1 and TCA. TCA was docked with a physics-based docking tool, AutoDock^57,58^, to identify its preferred coordination mode. These structures were further geometry optimized in large quantum mechanical (QM) cluster models using DFT. After the DFT optimization, the distance between the two Zn^2+^ ions was 3.5 Å, with the distance to the bridging hydroxide 1.9–2.0 Å for each Zn^2+^. While both Zn^2+^ were four-coordinate, the amide bond oxygen of TCA only coordinated α-Zn^2+^ (i.e., within 2 Å). This asymmetric coordination of α-Zn^2+^ was only observed in the DFT optimized structure, while docking tools placed the carbonyl oxygen at the midpoint of the two ions. In our model, the amide bond of TCA is positioned to undergo hydrolysis, and the negatively charged sulfonate group is recognized through hydrogen bonding and/or electrostatic interactions with Ser194, Gln381, Asn443, and His544 (**Fig. 4a**). A conserved histidine (His196 in *Bw*BSH1) is predicted to activate the amide carbonyl via hydrogen bonding and electrostatic interactions, supplementing the Lewis acid catalysis facilitated by coordination to the β-Zn^2+^ ion. Concurrently, a bound water molecule is activated for nucleophilic attack via interaction with the ɑ-Zn^2+^ ion.^38^ This dual polarization of the carbonyl group by a histidine and a β-metal ion is a recognized catalytic strategy within the amidohydrolase superfamily.^38,44,59^ Notably, sequence alignments revealed that the predicted sulfonate binding residues are conserved among *Bw*BSH homologs but are absent in other representative metal-dependent amidohydrolases (**Fig. 4b**). To test the predictions from our docking model, we evaluated the *Bw*BSH H196A mutant (**Fig. 4c**) and observed the complete loss of TCA hydrolysis activity.

**Fig. 4.**
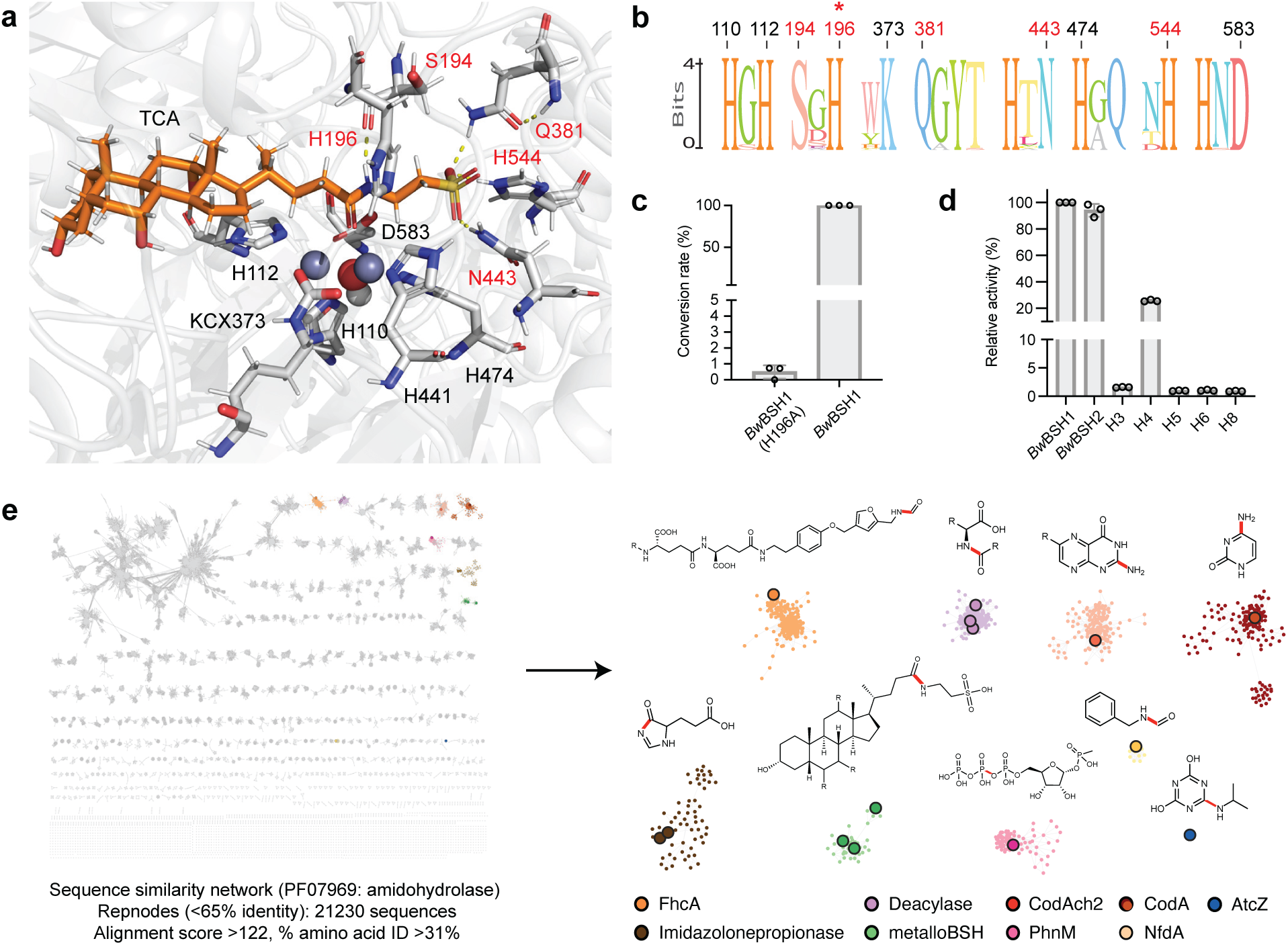
MetalloBSHs are distinct from other amidohydrolase superfamily members. a, AutoDock Vina model of TCA in the active site of AlphaFold3-predicted *Bw*BSH1 (containing a carboxylated lysine PTM, two Zn^2+^ ions, and a hydroxide). Ser194, Gln381, Asn443, and His544 (red) are predicted to stabilize the TCA sulfonate group via hydrogen bonding and/or electrostatic interactions. Metal-coordinating residues are labelled in black. The conserved His196 sits near the amide bond, consistent with an auxiliary role in Zn^2+^-dependent catalysis. b, Multiple sequence alignment highlighting conserved residues in metalloBSHs that may dictate specificity for taurine-conjugated BAs (substrate-binding residues in red; metal-binding residues in black). c, Complete loss of TCA hydrolysis in the H196A mutant supports a critical role for this residue in amide bond activation. d, TCA hydrolysis by short *Bw*BSH1-like amidohydrolases from *B. wadsworthia* 3_1_6. For panels c and d, bars represent the mean of three biological replicates, and error bars indicate the standard deviation. e, Sequence similarity network (SSN) of amidohydrolase family PF07969. Nodes representing uncharacterized sequences are shown in light gray. Nodes representing characterized sequences are distinctly colored, and experimentally validated enzymes are represented by enlarged nodes, including: imidazolonepropionase^60–62^, cytosine deaminase (CodA)^63,64^, pterin deaminase (CodAch2)^65^, aminoacylase^66–68^, formyltransferase/hydrolase (FhcA)^69^, *N*-substituted formamide deformylase (NfdA)^70^, *N*-isopropylammelide amidohydrolase (AtzC)^71^, and ⍺-D-ribose 1-methylphosphonate triphosphate (RPnTP) diphosphatase (PhnM)^72^.

Another feature that distinguishes *Bw*BSHs from most other metal-dependent amidohydrolases is their additional C-terminal domain. We probed the functional relevance of this domain by constructing a truncated *Bw*BSH1 variant (**Supplementary Table 6, Supplementary Fig. 11**). This construct exhibited a ∼4-fold loss of BSH activity in vitro (**Extended Data Fig. 4a, b**), indicating that while the N-terminal amidohydrolase domain is sufficient to support baseline BSH activity, the C-terminal domain is important for the catalytic efficiency of *Bw*BSHs. We also noted that, in addition to its two secreted *Bw*BSH homologs, *B. wadsworthia* 3_1_6 encodes 10 shorter amidohydrolases (482–572 amino acids, 23–32% aa sequence identity with *Bw*BSH1, named H2–H11) that lack the C-terminal domain (**Supplementary Table 7, Supplementary Fig. 12, Extended Data Fig. 4c**). Of these 10, only one (H8) is predicted to be secreted. Expression of these enzymes and activity assays revealed that while several (H3, H4, H5, H6, and H8) exhibit detectable BSH activity, their relative activity (**Fig. 4d**) and conversion rates (**Extended Data Fig. 4d**) were ∼4- to 100-fold lower, with H3 (WP_005026994) and H4 (WP_016360969) displaying the highest activity among this subset. These results highlight the C-terminal domain as a key feature impacting catalytic efficiency of *Bw*BSHs and suggest that additional amidohydrolases encoded in *B. wadsworthia* likely make more minimal contributions to BSH activity.

To further examine the relationship between metalloBSHs and other members of the metal-dependent amidohydrolase superfamily, we performed sequence similarity network (SSN) analysis.^73,74^ Although the vast majority of this superfamily remains uncharacterized, members with experimentally validated activities are represented by enlarged nodes. Using a minimum aa sequence identity of 31% and an alignment score cutoff of 122, our SSN resolves all characterized metal-dependent amidohydrolases with distinct activities into separate clusters (**Fig. 4e, Supplementary Table 8**). To verify that the *Bw*BSH-containing cluster (**Supplementary Table 9a**) contains bona fide metalloBSHs, we aligned the sequences using MAFFT^75^ to confirm the conservation of metal-binding ligands and the key active-site residues identified in our docking studies. This analysis allowed us to filter out partial or truncated sequences lacking a complete dinuclear metal center, yielding a curated set of 371 metalloBSH proteins (**Supplementary Table 9b**). A subcluster connected to the primary *Bw*BSH1 cluster contained additional enzymes with lower aa sequence similarity to *Bw*BSH1 (**Extended Data Fig. 5a**). To confirm that these divergent enzymes retain BSH activity, we heterologously expressed a representative candidate, *Tm*BSH1 (33.9% aa sequence identity to *Bw*BSH1), in *E. coli*. *Tm*BSH1 (WP_334315222.1) is encoded by *Taurinivorans muris*, a taurine-respiring mouse gut bacterium.^76^ Interestingly, while a prior study highlighted that *T. muris* does not encode Ntn-BSHs, its colonization in gnotobiotic mice increased the deconjugation of taurine-conjugated BAs^76^. Resolving this discrepancy, we found that purified recombinant *Tm*BSH1 hydrolyzed TCA with comparable efficiency to *Bw*BSH1 (**Extended Data Fig. 5b, 5c, Supplementary Table 10**). This further supports that this SSN cluster represents a family of metalloBSHs. Notably, the shorter amidohydrolases from *B. wadsworthia* that possess weak BSH activity segregate into a distinct uncharacterized cluster (**Extended Data Fig. 6**).

We next sought to characterize the taxonomic distribution of metalloBSHs. We analyzed representative genomes from the Genome Taxonomy Database (GTDB)^77,78^ and found that putative metalloBSHs are predominantly encoded within the *Desulfovibrionaceae* family (**Supplementary Table 11**). These organisms are members of the gut microbiomes of humans^79^, rodents^76^, porcine^80^, and avian species^81^, with some occasionally identified in clinical infections^82–85^. This phylogenetic profile suggests that while metalloBSHs may be largely restricted to *Desulfovibrionaceae*, they are widely distributed across diverse vertebrate microbiomes. This distribution highlights the likely importance of these enzymes for bacterial growth in bile-rich host environments.

To determine whether BSH activity is a core function in these bacteria, we examined the distribution of *Bw*BSHs across the *Bilophila* genus using the RefSeq genomes collection (as of 9/11/2025)^86^. We found that nearly all *Bilophila* strains encode *Bw*BSH and frequently encode multiple copies (**Supplementary Table 12**). Only 3 out of the 99 genomes assigned to the genus lack *Bw*BSH, and two of these are metagenome-assembled genomes and are thus likely incomplete. Resequencing of the other isolate that appeared to lack *Bw*BSH (DFI 5.69, GCF_020709195) revealed that it encodes three tandem-arranged *Bw*BSHs. These findings indicate that *Bw*BSHs are a highly conserved feature of *B. wadsworthia*, further suggesting they play a critical role in allowing these bacteria to inhabit the human gut.

#### *Metallobsh*s are expressed in vivo and prevalent in the human gut

We next sought to explore the relevance of metalloBSHs within the gut microbiome by searching for the sequences delineated by our SSN analysis in existing multi-omics datasets. First, we assessed their expression in a metatranscriptomic dataset from a study examining how *B. wadsworthia* ATCC 49260 exacerbates high-fat diet-induced metabolic dysfunction in Altered Schaedler Flora (ASF)-colonized mice. We found that all three *bwbsh* genes are among the top 10% most highly expressed protein-encoding genes in this strain in vivo (**Fig. 5a, Supplementary Table 13a**). Of the five short amidohydrolases exhibiting lower BSH activity, only *h3* is expressed at a level similar to *bwbsh*s. The top 1% of expressed genes includes key elements of taurine metabolism (*tpa*, *ald*, *islA*, and a gene encoding a TRAP transporter substrate binding protein), underscoring the likely importance of *Bw*BSHs in supplying taurine for growth.^87^ High *bwbsh* expression persisted even following the administration of the probiotic *Lactobacillus rhamnosus* CNCM I-3690 (**Fig. 5b, Supplementary Table 13b**). These data confirm the robust in vivo expression of genes encoding metalloBSHs.

**Fig. 5.**
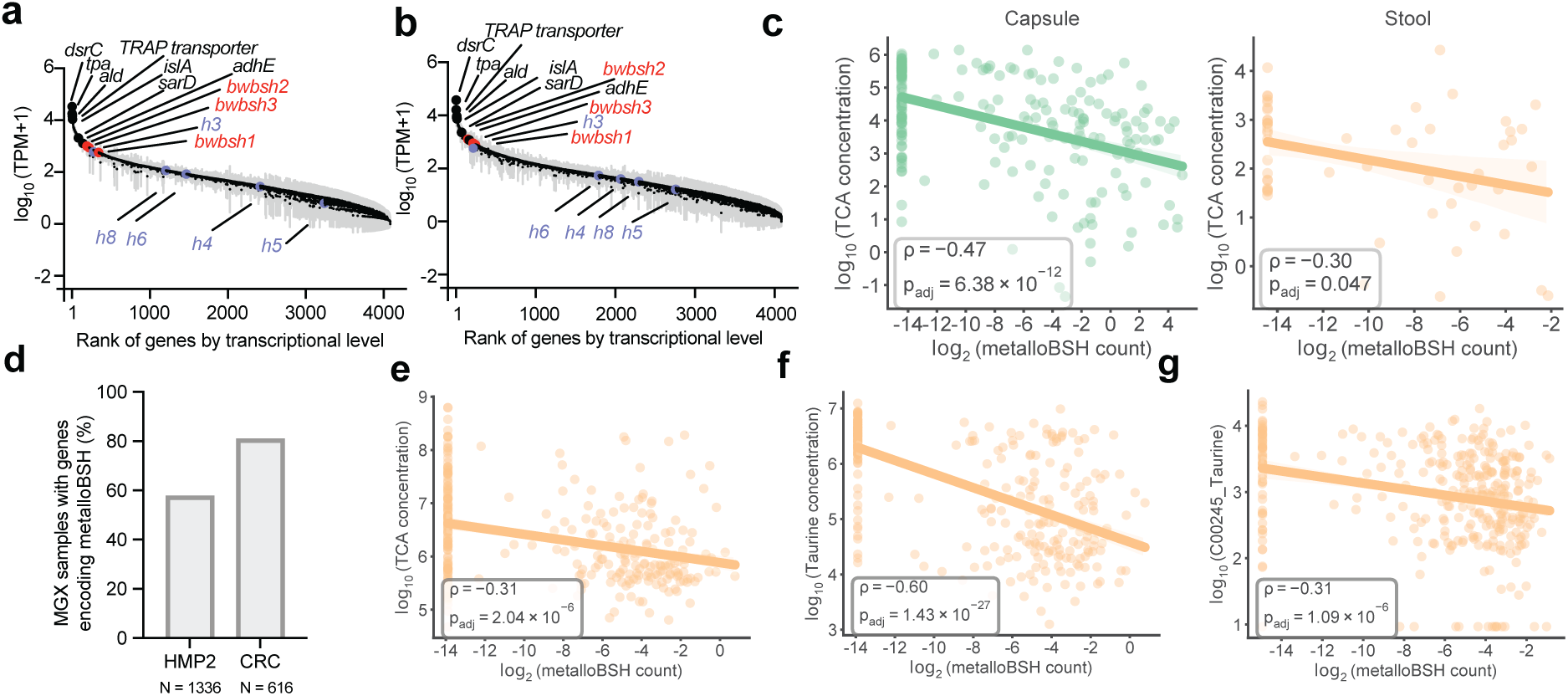
***Metallobshs* are present in the human gut microbiome and correlated with BA metabolism.** a, b, Ranked median transcriptional levels of the whole-protein-coding genes of *B. wadsworthia* ATCC 49260 in cecal samples of ASF-colonized mice in the absence (a) or presence (b) of *Lactobacillus rhamnosus* CNCM I-3690. Gray bar represents 95% confidence interval. TPM, transcripts per kilobase million. c, the log_2_ (read count) of metalloBSH-encoding genes in intestinal or stool metagenomes was negatively correlated with TCA levels in intestinal (*P_adj_* = 6.38 × 10^-12^) and stool (*P_adj_* = 0.047) metabolomes. Correlations are Spearman correlations with Benjamini–Hochberg correction. Data points are individual intestinal (N = 209) or stool samples (N = 56) for which both metagenome sequencing data and metabolomics data were available. d, the percentage of samples with metalloBSHs detected from MGX dataset of HMP2 cohort (N = 1336) and CRC cohort (N = 616). e, f, the log_2_ (read count) of metalloBSH-encoding genes in stool metagenomes (N = 297) from HMP2 dataset was negatively correlated with TCA (*P _adj_* = 2.04 ×10^-6^) and taurine levels (*P_adj_* = 1.43 ×10^-27^) in stool metabolomes. Correlations are Spearman correlations with Benjamini–Hochberg correction. Data points are individual stool samples (N = 297) for which both metagenome sequencing data and metabolomics data were available. g, log_2_ (read count) of metalloBSH-encoding genes in stool metagenomes from a CRC dataset was negatively correlated with taurine levels (*P_adj_* = 1.09 ×10^-6^) in stool metabolomes. Correlations are Spearman correlations with Benjamini–Hochberg correction. Points are individual stool samples (N = 347) for which both metagenome sequencing data and metabolomics data were available.

To examine the prevalence of metalloBSHs and their potential impacts on BA metabolism in vivo, we first analyzed a multi-omics dataset from 240 spatially resolved intestinal samples collected from humans via pH-responsive ingestible capsules.^29^ Although the original study correlated *B. wadsworthia* abundance with decreased levels of TCA in both intestinal samples and stool samples, consistent with BSH activity, it identified no corresponding BSH-encoding genes in this organism. Querying this dataset with the metalloBSHs from our SSN (**Supplementary Table 9b**) revealed that metalloBSH gene abundance was significantly negatively correlated with levels of all six taurine-conjugated bile acids analyzed in this study (**Fig. 5c**, **Extended Data Fig. 7**). These correlations were more pronounced in intestinal samples than in stool, likely because of the higher intestinal abundance of *Desulfovibrionaceae*. In contrast, we found weaker correlations between *metallobsh* abundance and levels of two of the six glycine-conjugated bile acids analyzed in both capsule and stool samples (**Extended Data Fig. 8**). Notably, analysis of *metallobsh* gene abundance in samples collected via localized devices at the pyloric sphincter (Capsule 1), small intestine (Capsule 2, Capsule 3), and ascending colon (Capsule 4) revealed a significant negative correlation with taurine-conjugated BAs specifically in the small intestine with the most significant negative correlations observed with TCA, TCDCA and TUDCA. No significant correlation (*P_adj_* > 0.05) was observed in the ascending colon samples (Capsule 4) (**Extended Data Fig. 9**). These inverse correlations are observed with three of the six detected glycine-conjugated BAs, albeit weaker than those observed for the corresponding taurine-conjugated BAs (**Supplementary Fig. 13**). Although the small intestine harbors a much lower microbial density than the colon, it is the site of ∼95% of BA reabsorption.^12^ Therefore, this robust correlation highlights a likely role for metalloBSH in modulating the systemic pool of taurine-conjugated BAs in the human body.

To further understand the distribution of metalloBSHs in the human gut and their impact on BA metabolism, we analyzed their prevalence in stool metagenomes from multiple clinical cohorts with matched stool metabolomics data, including studies focused on CRC^88^ and IBD^89^. *Metallobsh* genes are highly prevalent in the human gut microbiome: they were detected in 57%–80% of the metagenomes in the HMP2, 78%–89% of CRC cohort metagenomes, and ∼78% of individuals without diseases in both cohorts (**Fig. 5d**). Preliminary analyses also indicated that *metallobsh* prevalence correlates with disease across both cohorts (**Extended Data Fig. 10**).

Analyzing matched stool metabolomics data from the HMP2 cohort revealed *metallobsh* gene abundance is negatively correlated with levels of taurine-conjugated BAs (**Fig. 5e, Supplementary Fig. 14**) and taurine (**Fig. 5f**). Weaker correlations were observed with levels of glycine-conjugated BAs (**Supplementary Fig. 15**). In the data from the CRC cohort, a significant negative correlation was observed between *metallobsh* abundance and levels of taurine, but not TCA or GCA (**Fig. 5g**). Collectively, these findings demonstrate that *metallobshs* are prevalent in the mammalian gut microbiome and their abundance is associated with changes in levels of conjugated BAs and their downstream metabolites. Importantly, these data highlight metalloBSHs as active drivers of bile salt hydrolysis in vivo, revealing an overlooked dimension of BA metabolism by the gut microbiome.

## Discussion

BSH activity in the human gut microbiome has been a major focus of investigation due to its role as a gateway to further microbial modifications of BAs and its strong connections to host health and disease.^2,3^ Prior studies of BSHs have assumed this transformation is catalyzed by a single class of gut bacterial enzymes, the cysteine-dependent canonical Ntn-BSHs. Notably, efforts to characterize this activity in gut microbiome datasets and efforts to develop BSH inhibitors have focused solely on Ntn-BSHs, despite accumulating evidence linking organisms lacking this enzyme to BA metabolism.^29,31,90^ The discovery of metalloBSHs greatly expands our knowledge of this important gut microbial enzymatic activity.

While both classes of BSHs catalyze amide bond hydrolysis, biochemical characterization and structure prediction reveal important distinctions between metalloBSHs and Ntn-BSHs. Rather than employing a protein-based nucleophile, metalloBSHs likely use their metallocofactor to activate water for direct nucleophilic attack on conjugated BA substrates. This mechanistic distinction is reflected in the lack of susceptibility of metalloBSHs to covalent inhibitors of Ntn-BSHs and their inhibition by metal chelators. Multiple metal ions appear to support the activity of *Bw*BSH1, with maximal activity observed in the presence of Zn^2+^ and inhibition occurring with Cu^2+^. While additional studies are needed to precisely characterize the native metallocofactor of these enzymes, our findings align with studies of many other metal-dependent amidohydrolases^38,59^. Furthermore, metalloBSHs display high catalytic efficiency unmatched by most characterized Ntn-BSHs. This high activity, as well as the strict specificity of metalloBSHs for taurine-conjugated BAs and their extracellular localization, supports a potential role for these enzymes in accessing taurine to support the growth of *B. wadsworthia* and other taurine-respiring *Desulfovibrionaceae* in the gut.

Analyses of existing gut microbiome multi-omics datasets provides strong evidence for the relevance of metalloBSHs within mammalian gut microbiomes. They are highly expressed in vivo and are prevalent in datasets from clinical studies. Notably, the abundance of metalloBSH-encoding genes correlates with levels of conjugated BAs and BA metabolites in multiple cohorts. Furthermore, in our analysis of multi-omics data we find stronger correlations between the abundance of *metallobshs* and levels of BA metabolites compared to those previously reported for canonical Ntn-BSHs in intestinal datasets.^29^ Given that *B. wadsworthia* and BA metabolism have separately been connected to various health outcomes, our findings now set the stage for investigating the biological impacts of BSH activity in this organism.

Finally, our work further highlights the vast, untapped biochemical diversity that awaits characterization within large enzyme superfamilies, including transformations that are assumed to be thoroughly characterized.^91,92^ The discovery of metalloBSHs underscores the pitfalls of relying only on existing enzyme annotations in profiling metabolism in the gut microbiome, the value of activity-based enzyme approaches in uncovering gut microbial enzymes, and the possibility that additional important gut microbial transformations are catalyzed by a wider range of enzymes than is currently appreciated.

## Methods

### Materials and general methods

Primer synthesis and DNA sequencing were performed by Genewiz (Waltham, MA). Whole plasmid and whole genome sequencing were performed by Plasmidsaurus (Eugene, OR). DNA purification of recombinant plasmids was performed using an E.Z.N.A Plasmid DNA Mini Kit from Omega Bio-Tek (Norcross, GA). Restriction enzymes were purchased from New England Biolabs (Ipswich, MA), and digests were performed according to the manufacturer’s protocol. NEB Builder HiFi DNA Assembly Master Mix was purchased from New England Biolabs (Ipswich, MA) and used according to the manufacturer’s protocol. Nickel nitrilotriacetic acid agarose (Ni-NTA) resin was purchased from Qiagen (Germantown, MD) and ThermoFisher Scientific (Waltham, MA). Novex Tris-Glycine SDS-PAGE gels were purchased from ThermoFisher Scientific (Waltham, MA). Protein concentrations were determined using a Bradford assay (BioRad, CA). All chemicals were from Sigma-Aldrich unless otherwise noted. Taurine (*D*_4_) (Taurine-*d*_4_) and cholic acid (2, 2, 4, 4-*D*_4_) (CA-*d*_4_) were purchased from Cambridge Isotope Laboratories. Phusion/Q5 High-Fidelity PCR Master Mix was purchased from New England Biolabs (NEB). Hungate tubes were obtained from ChemGlass (Anaerobic Hungate Culture Tubes, GPI Screw Threads, 16 × 125 mm). Glycocholic acid (GCA) was purchased from Matrix Scientific. Taurodeoxycholic acid, sodium salt (TDCA) was purchased from AmBeed. Anaerobe basal broth (ABB) powder was obtained from HiMedia. Stock solutions of taurine, cholic acid, and bile salts were stored at –20 °C unless otherwise noted. All biological samples were stored at –80 °C when not in use unless otherwise noted. A personal H_2_S detector was used when exposing *B. wadsworthia* cultures to air to ensure that H_2_S exposure did not exceed safety thresholds.

*Bilophila wadsworthia* 3_1_6 and *Bilophila sp* 4_1_30 were obtained through BEI Resources (NIAID, NIH as part of the Human Microbiome Project). *Bilophila wadsworthia* ATCC 49260 was obtained from the American Type Culture Collection (ATCC). *Bilophila wadsworthia* DFI 5.69 was obtained from the Duchossois Family Institute Symbiotic Bacterial Strain Bank (DFI SBSB).

### *B. wadsworthia* culturing conditions

All culturing took place in a Coy Vinyl Chamber (97% N_2_/ 3% H_2_) equipped with a hydrogen sulfide removal column. Anaerobe basal broth (ABB) was prepared and autoclaved according to the manufacturer’s instructions. The freshly autoclaved medium was brought into the Coy Vinyl Chamber while still hot. Once cooled, 2 g/L sodium bicarbonate was added. Any additional sulfur, nitrate, or bile acids were added as solids where appropriate and then the medium was passed through a 0.22 μm syringe filter into sterile Hungate tubes in 5 mL aliquots.

For initial terminal electron acceptor screening, a single starter culture of *B. wadsworthia* 3_1_6 was inoculated from a glycerol stock into 5 mL of ABB + 5 mM taurine and allowed to grow at 37 °C until saturated. ABB media was prepared in 4 × 5 mL portions per additive (5 mM isethionate, GCA, nitrate, sulfite, taurine, TCA, or TDCA). One aliquot was sealed to serve as a media blank for each condition, and the other three were inoculated with 25 μL of the saturated starter culture. The cultures were sealed and removed from the Vinyl Chamber and OD_600_ readings were taken using a GENESYS 20 Visible Spectrophotometer (Thermo Scientific). Each media blank was used to zero the spectrophotometer. When not being measured, cultures were stored at 37 °C wrapped in aluminum foil. Timepoints were measured as indicated from 0–353.75 hours. All readings OD_600_ > 2.5 are outside the listed photometric range but are included for comparison purposes. When cultures reached an OD_600_ = 4, the instrument reached a maximum and growth measurements were stopped.

For comparing growth across strains (*B. wadsworthia* 3_1_6, *B. wadsworthia* ATCC 49260 and *Bilophila* sp. 4_1_30), each strain was inoculated from a glycerol stock into 5 mL of ABB + 5 mM taurine medium. All starter cultures were grown for the same amount of time (3 days). ABB media was prepared with either 5 mM taurine, 5 mM TCA or 5 mM GCA and filtered and aliquoted into 5 mL volumes in Hungate tubes. One aliquot for each additive was sealed as a media blank, and the remaining tubes were inoculated in triplicate per media condition and with 25 μL of one of the three starter cultures. Cultures were sealed and brought out of the anaerobic chamber to take OD_600_ readings using the GENESYS 20 Visible Spectrophotometer. When not being measured, cultures were stored at 37 °C and wrapped in aluminum foil. Timepoints were recorded over 0–136.5 hours.

### Bile salt deconjugation during *B. wadsworthia* growth

A single 5 mL culture of ABB + 5 mM sodium sulfite in a Hungate tube was inoculated from a *B. wadsworthia* 3_1_6 glycerol stock. The culture was allowed to grow at 37 °C until saturated. ABB + 5 mM TCA medium was prepared and aliquoted into 5 mL portions in Hungate tubes. Media blanks were sealed and the experimental cultures were inoculated with 50 μL of the saturated starter culture. Cultures were prepared in triplicate. The sealed cultures were removed from the anaerobic chamber and OD_600_ readings were taken in a GENESYS 20 Visible Spectrophotometer. Immediately following OD_600_ measurements, a sterile 100 μL aliquot was withdrawn with a needle through the rubber septum of each culture. 10 μL of the culture aliquot was added to 90 μL of acetonitrile (ACN) and mixed. Samples were stored at –70 °C until UHPLC-MS/MS analysis. Cultures were returned to 37 °C and wrapped in aluminum foil. Timepoints were taken at between 0–45.75 hours post-inoculation. For UPLC-MS/MS analysis, ACN extracts were thawed and centrifuged at 16,000 × *g* for 15 min at 4 °C. 4 μL of the supernatant was further diluted into 196 μL of ACN in a clear 96-well plate. Standard curves of TCA, CA, and taurine were prepared in a combined standard curve. 10 μL of standards were added to 90 μL of ACN and processed similarly to the samples. Standards were run in triplicate.

#### UPLC-MS/MS analytical methods

UHPLC-MS/MS analysis was performed on a Waters Xevo TQ-S system equipped with a triple quadrupole and an electrospray ionization (ESI) source in negative mode. A Waters Acquity BEH/Amide UPLC column (1.7 μm, 130 Å, 2.1 × 50 mm, heated to 40 °C) was injected with 1 μL of sample at a column flow rate of 0.5 mL min^-1^. The LC conditions were as follows using solvent A (H_2_O with 0.1 % formic acid) and solvent B (ACN with 0.1 % formic acid): 0–0.3 min 97–73% B, 0.3–1 min 73% B, 1.0–1.5 min 73–30% B, 1.5–2 min 30% B, 2–2.2 min 30–97% B, 2.2–4 min 97% B; column flow was directed to the MS/MS from 1.1–2.5 min. ESI negative mode was used to measure TCA, CA, and taurine. Metabolite quantification parameters are listed in **Supplementary Table 14**. Transitions were optimized using the Waters Intellistart software. Concentrations of TCA, CA, and taurine were determined using TargetLynx.

#### Q-TOF analytical methods

An Agilent Q-TOF 6530 equipped with a Dual AJS ESI source was used for LC–HRMS analysis. A Cogent Diamond Hydride column (4 µm, 130 Å, 3 × 150 mm, Mirosolv Technology Corp) was injected with 1 μL of sample at a flow rate of 0.5 mL min^−1^. The LC conditions were as follows using solvent A (H_2_O with 0.1 % formic acid) and solvent B (ACN with 0.1 % formic acid): 0–2.0 min 97–73% B, 2.0–7.0 min 73% B, 7.0–10.0 min 73–30% B, 10.0–15.0 min 30% B, 15.0–16.0 min 30–97% B, 16.0–21.0 min 97% B; column flow was directed to the HRMS from 2.0–15.0 min. The following parameters were used for the Q-TOF: Gas Temp 300 °C, Drying Gas 8 L/min, Nebulizer 35 psi, Sheath Gas Temp 350 °C, Sheath Gas Flow 11 L/min, VCap 3500 V, Nozzle Voltage 1000 V.

### Bile salt deconjugation in resuspended *B. wadsworthia* cultures

A single 5 mL culture of ABB + 5 mM taurine in a Hungate tube was inoculated from a *B. wadsworthia* 3_1_6 glycerol stock. The culture was allowed to grow at 37 °C until saturated. ABB + 5 mM taurine medium was prepared and aliquoted in 5 mL portions into Hungate tubes. 2 × 5 mL cultures were inoculated with 25 μL of starter culture, sealed, and grown at 37 °C overnight. The two cultures reached an average OD_600_ of 1.32 as measured using a GENESYS 20 Visible Spectrophotometer. All subsequent steps took place aerobically in a fume hood. The two cultures were combined to attain higher cell amounts and centrifuged at 4,000 × *g* for 4 min at room temperature. The supernatant was decanted and the pellet was resuspended in phosphate buffered saline (PBS, 137 mM NaCl, 2.7 mM KCl, 10 mM Na_2_HPO_4_, 1.8 mM KH_2_PO_4_, pH 7.4). The cells were centrifuged again and the supernatant was decanted. This wash step was repeated twice. The final cell pellet was resuspended in 750 μL of PBS. 325 μL of resuspended cells were heated to 80 °C for 10 min as a heat-treated control. 5 mM stocks of TCA, TDCA, and GCA were prepared in PBS. 32.5 μL of resuspended cells or blank PBS were mixed with 32.5 μL of bile salt stock solutions to reach a working concentration of 2.5 mM. Each condition and bile salt mix was prepared in triplicate. Timepoints were taken as indicated over a 90-minute incubation. At each timepoint, 10 μL of assay mixture was diluted into 90 μL of ACN containing CA-*d*_4_ and Taurine-*d*_4_ as internal standards. ACN extracts were stored at –20 °C to await same day sample processing. Extracts were thawed and centrifuged at 16,000 × *g* for 15 min at 4 °C. 1.2 μL of supernatant was diluted into 58.8 μL of ACN in a 384-well plate that was sealed with a pierceable plate seal. Two sets of mixed standards were prepared (TCA, CA, and taurine; TDCA and GCA) and were processed similarly to the samples. Standards were run in duplicate. UPLC-MS/MS analysis was performed as described in **UPLC-MS/MS analytical methods**. Concentrations of TCA, TDCA, GCA, CA, and taurine were determined using Target Lynx. CA-*d*_4_ was used as an internal standard for CA, TCA, TDCA, and GCA quantification. Taurine-*d*_4_ was used as an internal standard for taurine quantification.

#### Testing the inducibility of *Bw*BSH

A single 5 mL culture of ABB + 5 mM taurine in a Hungate tube was inoculated from a *B. wadsworthia* 3_1_6 glycerol stock. The culture was allowed to grow at 37 °C until saturated. ABB + 5 mM sulfite medium was prepared and aliquoted in 5 mL portions into Hungate tubes. 4 × 5 mL cultures were inoculated with 25 μL of starter culture, sealed, and grown at 37 °C overnight. The two cultures with the most similar OD_600_ readings as measured in the GENESYS 20 Visible Spectrophotometer were chosen for further studies. One culture was supplemented with 200 μL of 25 mM TCA in anoxic PBS (OD_600_ = 1.31), while the other culture was supplemented with 200 μL of anoxic PBS (OD_600_ = 1.36), both via needle through the septum to minimally disrupt the headspace. The cultures were allowed to grow for an additional 90 min at 37 °C wrapped in foil. The two cultures had similar ODs post-incubation (OD_600_ = 1.530 and 1.501 for PBS blank and TCA supplemented, respectively). All subsequent steps took place aerobically in a fume hood. Each culture was centrifuged at 4,000 × *g* for 4 min at room temperature. The supernatant was decanted and the pellets were resuspended in PBS. This wash step was repeated twice. The final cell pellets were each resuspended in 500 μL of PBS. A 5 mM stock of TCA was prepared in PBS. 75 μL of resuspended cells or PBS were mixed with 75 μL of TCA stock solution to reach a working concentration of 2.5 mM. Each culture resuspension was prepared in triplicate, and the corresponding PBS control incubations were prepared in duplicate. Timepoints were taken as indicated over a 60-minute incubation. At each time point, 10 μL of assay mixture was diluted into 90 uL of ACN containing CA-*d*_4_ and Taurine-*d*_4_ as internal standards. ACN extracts were entrifuged at 16,000 × *g* for 15 min at 4 °C. 4 μL of supernatant was diluted into 196 μL of ACN in a 96-well plate that was sealed with a pierceable plate seal. UPLC-MS/MS analysis was performed as described in **UPLC-MS/MS analytical methods**. CA-*d*_4_ was used as an internal standard for CA and TCA quantification. Taurine-*d*_4_ was included to standardize taurine concentration.

#### Activity-guided purification of *Bw*BSH

*Preparation of cell free extract: B. wadsworthia* 3_1_6 cell pellets from a total of 2 L of culture were thawed and resuspended in a total of 45 mL of fresh lysis buffer (50 mM MOPS, 0.1 mg mL^-1^ DNase, 0.5 mg mL^-1^ lysozyme, pH 7.4) with added EDTA-free protease inhibitor. The suspension was sonicated twice at 25% amplitude, 2 s on, 8 s off, and 1 min total sonication time using a Sonifier 250 horn fitted with a 1/2” tip (Branson, Dansbury, CT). In between the sonication steps, the suspension was manually inverted several times. The lysate was clarified by centrifugation at 18,000 × *g* at 4 °C for 45 min.

*Anion-Exchange Chromatography (AEC):* The supernatant was loaded onto the AEC column (Cytiva HiTrap Q FF, 5 mL), which was equilibrated with a buffer of 50 mM MOPS buffer, pH 7.4 (Buffer A). The protein solution was loaded at 2.5 mL min^-1^ followed by an additional 5 mL of Buffer A. Protein was eluted with the same buffer using a linear gradient of NaCl from 0 to 500 mM from 100% at 0 mL to 0% at 705 mL of Buffer A at 2.5 mL min^-1^. The absorption of column effluents was monitored at 280 nm. 144 × 5 mL fractions were collected in 8 mL tubes throughout the entire run. AEC fractions were assayed for TCA deconjugation activity at five-fraction intervals. *Hydrophobic Interaction Chromatography (HIC):* Active samples, along with any intermediate fractions located between two active test intervals, were pooled and adjusted with a sample buffer to achieve a final concentration of 1000 mM ammonium sulfate prior to loading onto the HIC column (Cytiva HiTrap Phenyl HP, 5 mL), which was equilibrated with buffer containing 50 mM MOPS, 1000 mM ammonium sulfate buffer at pH 7.4 (Buffer B). The protein solution was loaded at 2.5 mL min^-1^ followed by an additional 5 mL of Buffer B. Protein was eluted using a linear gradient of ammonium sulfate from 1000 mM to 0 mM at 2.5 mL min^-1^. The absorption of column effluents was monitored at 280 nm. 144 × 5 mL Fractions were collected in 8 mL tubes throughout the entire run. AEC fractions were assayed for TCA deconjugation activity at five-fraction intervals. *Size-Exclusion Chromatography (SEC):* Active samples, along with any intermediate fractions located between two active test intervals, were pooled and concentrated to ∼130 μL before being subjected to SEC. The concentrated solution was loaded onto the SEC column (Superdex 200 Increase 10/300 GL), which was equilibrated with a buffer of 50 mM MOPS, 500 mM ammonium sulfate buffer at pH 7.4 (Buffer C). The protein solution was loaded at 0.375 mL min^-1^ followed by an additional 5 mL of Buffer C. Protein was eluted with an isocratic flow of Buffer C at 0.375 mL min^-1^ for a total volume of 1.3 Column Volumes (CVs). Absorption of column effluents was monitored at 280 nm. 20 × 1.3 mL fractions were collected in a 96-well deep well plate from approximately 6.2 mL to 32.2 mL during the SEC run. Fourteen fractions from approximately 6.2 mL to 24.4 mL were tested for activity.

Due to the large number of AEC and HIC fractions collected, one of every five AEC/HIC fractions were tested for TCA deconjugation activity, in addition to the eight AEC and eighteen HIC flowthrough fractions. All fractions assessed were tested as single replicates. 25 μL of each selected fraction was mixed 1:1 with 5 mM TCA in PBS, with time points at 0 min and 60 min of incubation time at 37 °C. At each time point, 10 μL of the assay mixture was diluted into 90 μL of ACN containing 11.1 μM Taurine-*d*_4_ and 11.1 μM CA-*d*_4_ as internal standards. For UPLC-MS/MS analysis, ACN extracts were centrifuged at 16,000 × *g* for 15 min at 4 °C. 4 μL of the supernatant was further diluted into 196 μL of ACN in a clear 96-well plate and sealed with a pierceable plate seal. Standard curves of TCA, CA, and taurine were prepared separately, and 10 μL of standards were added to 90 μL ACN with internal standards and processed similarly to the samples. UPLC-MS/MS analysis was performed as above with the transitions monitored listed in **Supplementary Table 14**. CA-*d*_4_ was used as an internal standard for CA and TCA quantification. Taurine-*d*_4_ was included to standardize taurine concentration.

#### Protein sequence analysis by LC-MS/MS

Protein identification was performed by the Taplin Biological Mass Spectrometry Facility at Harvard Medical School (Boston, MA). Fractions exhibiting BSH activity were concentrated using Amicon Ultra centrifugal filters (3 kDa MWCO, MilliporeSigma) and resolved on a Novex 4–15% Tris–Glycine mini protein gel. Gel bands of interest were excised and cut into ∼1 mm³ pieces, followed by in-gel trypsin digestion.^93^ Briefly, gel pieces were washed and dehydrated with ACN for 10 min and then dried completely in a SpeedVac. Gel pieces were rehydrated at 4 °C in 50 mM ammonium bicarbonate containing 12.5 ng µL^-1^ sequencing-grade modified trypsin (Promega, Madison, WI). After 45 min, excess trypsin solution was removed and replaced with 50 mM ammonium bicarbonate to cover the gel pieces, and samples were incubated overnight at 37 °C. Peptides were extracted by removing the ammonium bicarbonate buffer and washing once with 50% ACN containing 1% formic acid. Combined extracts were dried in a SpeedVac (∼1 h) and stored at 4 °C until analysis.

For LC–MS/MS analysis, samples were reconstituted in 5–10 µL of HPLC solvent A (2.5% ACN, 0.1% formic acid). A nano-scale reverse-phase HPLC capillary column was created by packing 2.6 µm C18 spherical silica beads into a fused silica capillary (100 µm inner diameter × ∼30 cm length) with a flame-drawn tip. ^94^ After column equilibration, samples were loaded via a Famos autosampler (LC Packings, San Francisco, CA). Peptides were separated by an increasing gradient of solvent B (97.5% ACN, 0.1% formic acid). Eluting peptides were ionized by electrospray and analyzed on a Velos Orbitrap Pro ion-trap mass spectrometer (Thermo Fisher Scientific, Waltham, MA). Peptides were detected, isolated, and fragmented to generate tandem mass spectra. Protein identification was achieved by searching the spectra against protein databases using SEQUEST (Thermo Fisher Scientific).^95^ Databases included a reversed (decoy) version of the *Bilophila wadsworthia* proteome, and search results were filtered to achieve a peptide false discovery rate between 1–2%.

#### Cloning for protein expression

Forward and reverse primers (**Supplementary Table 15**) for all ten BSH candidates (WP_005024313, WP_005025135, WP_005025869, WP_005028227, WP_005028462, WP_005028464, WP_005028753, WP_005028821, WP_005029059, and WP_005030331), *Bw*BSH homologs (WP_016360761, WP_005024391, WP_005026994, WP_016360969, WP_016360533, WP_016360860, WP_016360918, WP_016360866, WP_005027200, WP_005030026, WP_009368663), and truncated *Bw*BSH1 were designed using the NEBuilder Assembly Tool. Genes were amplified from genomic DNA using Q5 High-Fidelity 2 × Master Mix according to the manufacturer’s protocol (NEB). Due to the size of the construct for *Bw*BSH1 and *Bw*BSH2, two sets of primers were created that generated two fragments that could then be assembled to form the full construct. NdeI and XhoI restriction enzymes were used to obtain the backbone fragment of the pET28a(+) cloning vector. The gel-purified PCR products of respective BSH candidates and NdeI-XhoI digested backbone fragment of the pET28a(+) cloning vector were assembled using NEBuilder HiFi DNA Assembly Master Mix according to manufacturer instructions. 2 µL of each construct was used to transform 50 µL of chemically competent *E. coli* TOP10 cells. The cells were plated on LB supplemented with kanamycin (50 µg mL^-1^). The plasmids were isolated using an E.Z.N.A Plasmid Mini Kit I (Omega Bio-tek) and the identities of each of the resulting plasmids were confirmed by Sanger sequencing (Genewiz) or whole plasmid sequencing (Plasmidsaurus).

#### Site-directed mutagenesis

Site-directed mutagenesis of the gene encoding *Bw*BSH1 was achieved by using overlapping primers (**Supplementary Table 15**) to introduce mutations. Two fragments created from two sets of primers were assembled with NdeI-XhoI digested backbone fragment of the pET28a(+) cloning vector using NEBuilder HiFi assembly Master Mix. 2 µL of each construct was used to transform 50 µL of chemically competent *E. coli* TOP10 cells. The cells were plated on LB medium supplemented with kanamycin (50 µg mL^-1^). The plasmids were isolated using an E.Z.N.A Plasmid Mini Kit I (Omega Bio-tek) and the identities of each of the resulting plasmids were confirmed by whole plasmid sequencing (Plasmidsaurus).

#### Recombinant protein production and purification

1. **BSH candidates**

Plasmids pET28a(+) harboring the respective gene inserts were transformed into *E. coli* BL21(DE3) via heat shock, and the transformed strains were grown in LB supplemented with 50 μg mL^-1^ of kanamycin. After inoculation of 1 L of LB broth containing 35 μg mL^−1^ of kanamycin,

the cultures were grown at 37 °C until the cell density reached an A600 ≈ 0.6, when expression was induced with 0.15 mM isopropyl β-D-1-thiogalactopyranoside (IPTG). After induction, the cultures were grown at 16 °C for 20 h in a shaking incubator. Cell pellets were then harvested by centrifugation for 15 min at 5,000 × *g* at 4 °C. The cell suspension was mixed by pipetting and lysed via sonication for a total of 10 min at 25% amplitude with 2 s pulses separated by 8 s rest periods using a Sonifier S-250 horn fitted with a 1/2” tip (Branson, Dansbury, CT). The resulting lysate was clarified by centrifugation at 18,000 × *g* for 45 min. After centrifugation, the protein was purified using affinity chromatography with HisPur Ni-NTA agarose (Qiagen), and proteins were eluted with increasing concentrations of imidazole (20 mM–200 mM) in buffer A (50 mM HEPES, 10 mM imidazole, 500 mM NaCl, pH 8.0). Purified proteins were concentrated using Amicon Ultra centrifugal filters (50 kDa MW cutoff for WP_005025869, 10 kDa MW cutoff for WP_005029059, and 30 kDa MW cutoff for all other candidates); the concentrated solution was buffer exchanged using a PD-10 column (Cytiva) into storage buffer (20 mM HEPES, 50 mM NaCl, 10% glycerol, pH 8.0). Protein purity was assessed using Bio-Rad 4–15% gradient SDS-PAGE gel electrophoresis. Bio-Rad Precision Plus Protein™ All Blue Prestained Protein Standards was used as the ladder. His_6_-tagged proteins were used without further modifications.

1. *Bw*BSH1 and *Bw*BSH2

*Bw*BSH1 lacking its secretion signal peptide, and *Bw*BSH2 retaining its native signal peptide, were expressed in *E. coli* ArcticExpress (DE3) (Agilent Technologies) using the pET28a(+) expression vector. A frozen glycerol stock harboring the expression vector was used to inoculate an overnight culture in LB medium supplemented with 50 μg mL^-1^ kanamycin and 20 μg mL^-1^ gentamycin. The culture was grown at 37 °C with shaking at 200 rpm. For protein expression, 10 mL of the overnight culture was used to inoculate each liter of Terrific Broth (TB) containing 35 μg mL^-1^ kanamycin and 14 μg mL^-1^ gentamycin. Cultures were incubated at 37 °C with shaking at 200 rpm until reaching an OD_600_ of 0.5–0.7. Protein expression was induced by adding 150 μM IPTG, followed by incubation at 13 °C for 16 h with continued shaking at 180 rpm. Cells were harvested by centrifugation at 6,000 × *g* for 10 min and resuspended in lysis buffer (50 mM HEPES, 400 mM NaCl, 10 mM imidazole, pH 8.0). Cell lysis was performed using an Avestin EmulsiFlex C3 high-pressure homogenizer (15,000–20,000 psi). The lysate was clarified by centrifugation at 18,000 × *g* for 45 min, and the supernatant was applied to a Ni-NTA resin column (Qiagen/Thermo Fisher Scientific) pre-equilibrated in lysis buffer. The resin was washed with wash buffer (50 mM HEPES, 400 mM NaCl and 20 mM imidazole, pH 8.0) before eluting the protein of interest with elution buffer (50 mM HEPES, 400 mM NaCl, and 200 mM imidazole, pH 8.0). The protein was buffer-exchanged into storage buffer (20 mM HEPES, 100 mM NaCl, pH 8.0) using FPLC, flash-frozen in liquid nitrogen, and stored at −70 °C until use.

#### Activity assay with purified BSH candidates

Reaction mixtures consisted of PBS (pH 7.4), 1 mM TCA, and 1 μM enzyme at 37 °C in a final reaction volume of 100 μL. After 1 h, 10 μL of the reaction mixture was quenched by adding 90 μL of ACN followed by centrifugation (16,000 × *g* for 15 min) to remove the precipitated protein. The reaction was monitored by LC–MS as described in **UPLC-MS/MS analytical methods**.

#### General procedure for BSH activity assays

To evaluate BSH activity, 100 μL reactions were carried out in PBS (pH 7.4) at 37 °C, utilizing a final enzyme concentration of 5 nM and 1 mM of TCA (or alternative taurine-conjugated bile acids). The reactions were sampled at 15 min, 30 min, and 60 min by withdrawing 10 μL aliquots, which were subsequently quenched by the addition of 90 μL of ACN. Following centrifugation at 16,000 × *g* for 15 min to remove precipitated protein, the cleared supernatants were analyzed via established **UPLC-MS/MS analytical methods** for quantitative measurements **or Q-TOF MS analytical methods** for qualitative characterization.

**Conversion rate (%)** was expressed as the percentage of taurine product generated by each variant compared to wild-type *Bw*BSH1 following an overnight incubation.

**Relative activity (%)** was determined by normalizing the concentration of taurine product generated in enzyme-containing test group to that in the control group under initial rate conditions. **Kinetic analyses of *Bw*BSH1 and *Bw*BSH2**

Reaction mixtures consisted of PBS (pH 7.4) containing variable amounts of TCA (10–20000 μM). The reaction was performed at 37 °C with 5 nM *Bw*BSH1 or *Bw*BSH2, and samples were collected at multiple time points. The 15-min time point was selected as the analytical time point to ensure the reaction under initial velocity conditions (<10% conversion) over this time frame. Product formation was determined as described in **UPLC-MS/MS analytical methods**. Each data point represents a minimum of three replicate end-point assays; kinetic parameters were calculated by fitting the data to the Michaelis-Menten equation via nonlinear regression using GraphPad Prism v10.2.0.

#### pH dependence of *Bw*BSH1

To determine the pH dependence of *Bw*BSH1, enzymatic activity was measured across a pH range of 4.0 to 8.0. Assays were conducted using sodium acetate buffer (pH 4.0, 4.4, 5.0, 5.4) and sodium phosphate buffer (pH 6.0, 6.4, 7.0, 7.4, 8.0). All reaction mixtures contained a final concentration of 5 nM *Bw*BSH1 and 1 mM TCA. The reaction was monitored by UPLC-MS/MS as described in **UPLC-MS/MS analytical methods**.

#### Preparation of metal-depleted *Bw*BSH1 and metal reconstitution

Metal-depleted *Bw*BSH1 was prepared by two methods. Method 1: *Bw*BSH1 was expressed by cultures grown in M9 minimal medium supplemented with 0.2% casamino acids, 0.4% glucose, 2 mM MgSO_4_, and 0.1 mM CaCl_2_ and 35 µg mL^-1^ kanamycin. The purification of metal depleted *Bw*BSH1(M9) was as described in ***Bw*BSH1 and *Bw*BSH2 purification for assays**. Method 2: Purified *Bw*BSH1 expressed by cultures grown in Terrific Broth was purified and incubated with 5 mM 8-hydroxyquinoline-5-sulfonic acid and 5 mM 1,10-phenanthroline overnight in storage buffer. The metal-bound 8-hydroxyquinoline-5-sulfonic acid or metal-bound 1,10-phenanthroline was removed from *Bw*BSH1 via dialysis against PBS buffer (pH 7.4) for 24 h with three buffer changes, yielding the preparation referred to as *Bw*BSH1(chelators). The Zn(II)-reconstituted *Bw*BSH1 (chelators) was prepared by incubating 5 µM metal-depleted *Bw*BSH1(chelators) with 100 µM ZnCl_2_ in PBS buffer (pH 7.4) overnight and protein was separated from free Zn(II) or Zn(II)-reconstituted *Bw*BSH1 by passage through a PD-10 column equilibrated with PBS buffer(pH 7.4). Cu(II)-reconstituted protein was prepared using an identical procedure with CuCl₂. *Bw*BSH1 was expressed in M9 minimal medium supplemented with 0.2% casamino acids, 0.4% glucose, 2 mM MgSO4, 0.1 mM CaCl_2_, and 100 µM ZnCl_2_ to obtain in vivo Zn(II)-loaded *Bw*BSH1 (M9 + Zn(II)). **ICP-MS analysis of *Bw*BSH1**

Trace-metal-free nitric acid (60 µL) was added to 200 µL of an aqueous *Bw*BSH1 solution (2 mg mL^-1^). The mixture was incubated for 3 h at 60 °C, after which precipitates were removed by centrifugation. A 200 µL aliquot of the supernatant was diluted to a final volume of 3 mL with molecular biology grade water (Thermo Fisher Scientific). Samples were analyzed using an Agilent 7900 Inductively Coupled Plasma Mass Spectrometer (ICP-MS) at the MIT Center for Environmental Health Sciences (CEHS) Bio Core Facility. The isotopes selected for analysis were ^55^Mn, ^56^Fe, ^59^Co, ^60^Ni, ^63^Cu, and ^66^Zn. Instrument performance was optimized daily through autotuning followed by verification via a performance report.

#### Activity assay with metal-depleted *Bw*BSH1

Activity of recombinant *Bw*BSH1(LB), metal-depleted *Bw*BSH1(chelators), and metal-depleted *Bw*BSH1(M9) were tested by incubating 8 nM enzyme in 100 µL of PBS buffer containing either 24 nM ZnCl_2_, CuCl_2_, FeSO_4_, MnSO_4_, CoCl_2_, or NiCl_2_ (pH 7.4) for 30 min at 37 °C. TCA (1 mM) was added to the above solution and incubated at 37 °C for 15 min. A 10 μL aliquot of the reaction mixture was then quenched with 90 μL of ACN and centrifuged at 16,000 × *g* for 15 min to remove the precipitated protein. The reaction was monitored by UPLC-MS/MS as described in **UPLC-MS/MS analytical methods**.

#### Synthesis of cholylhydroxamic acid (CHA)

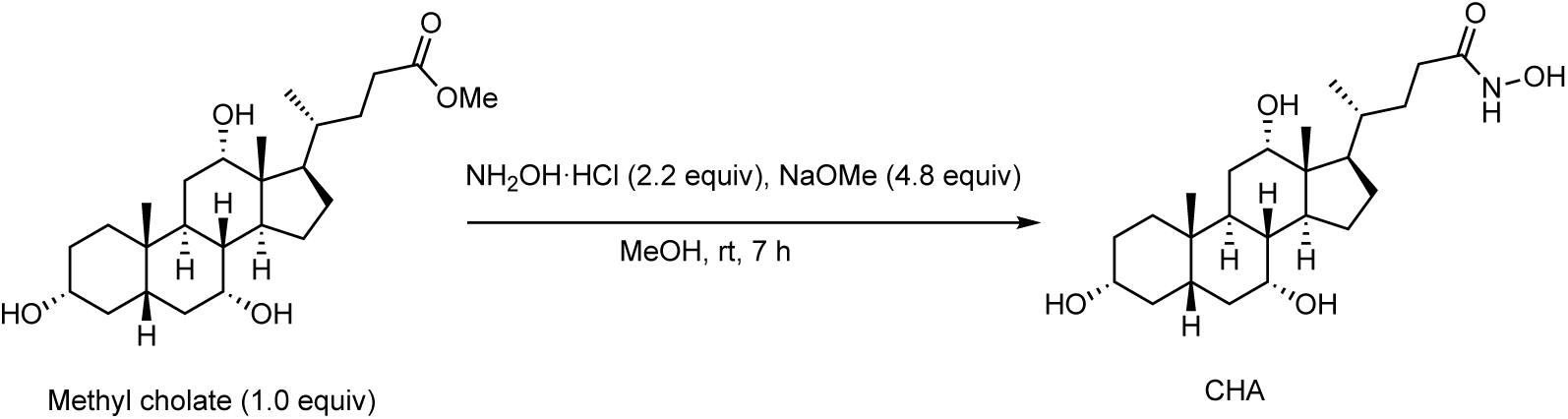

Hydroxylamine hydrochloride (NH_2_OH·HCl, 0.7 g, 10 mmol, 2.2 equiv) was dissolved in 1 mL of methanol and added to a solution of sodium methoxide (CH_3_ONa, 25 wt % in methanol, 5.07 mL, 22 mmol, 4.8 equiv).^50^ The resulting solution was added to a solution of methyl cholate (1.95 g, 4.6 mmol, 1.0 equiv) in 12 mL of methanol, and the resultant mixture was stirred at room temperature for 7 h.^51^ The reaction mixture was neutralized to pH 6.8 with hydrogen chloride solution (3 M in methanol), filtered, and concentrated under reduced pressure at 42 °C. Purification using reversed phase chromatography on a Biotage Selekt using a 6 g C18 column with a 10 to 90% gradient of ACN in water afforded cholylhydroxamic acid (1.78 g, 4.2 mmol, 91% yield) as white powder.

^1^H NMR (400 MHz, *CD*_3_*OD*) (**Supplementary Fig. 16**): δ 3.95 (t, *J* = 2.9 Hz, 1H), 3.80 (q, *J* = 3.0 Hz, 1H), 3.37 (tt, *J* = 11.1, 4.3 Hz, 1H), 2.35 – 2.21 (m, 2H), 2.16 (ddd, *J* = 13.4, 9.9, 4.9 Hz, 1H), 2.06 – 1.72 (m, 8H), 1.69 – 1.25 (m, 11H), 1.11 (qd, *J* = 11.8, 5.2 Hz, 1H), 1.03 (d, *J* = 6.4 Hz, 3H), 0.97 (dd, *J* = 14.1, 3.4 Hz, 1H), 0.92 (s, 3H), 0.71 (s, 3H). ^13^C NMR (101 MHz, *CD*_3_*OD*) (**Supplementary Fig. 17**): δ 173.58, 74.04, 72.88, 69.03, 47.96, 47.48, 43.20, 43.00, 41.01, 40.47, 36.83, 36.49, 35.90, 35.87, 33.09, 31.18, 30.76, 29.58, 28.66, 27.88, 24.22, 23.16, 17.65, 12.98.

HRMS (ESI+, *m/z*) (**Supplementary Fig. 18**): [M+H]^+^ calculated for C_24_H_42_NO_5_^+^, 424.3057; found 424.3044. [2M+H]^+^ calculated for C_48_H_83_N_2_O_10_^+^, 847.6042; found 847.6017. [2M+Na]^+^ calculated for C_48_H_82_N_2_NaO_10_^+^, 869.5862; found 869.5837.

#### Activity of recombinant *Bw*BSH1 with pan-BSH inhibitor BSH-IN-1 and CHA

To determine the effect of pan-BSH inhibitor on *Bw*BSH1 activity, 8 nM recombinant *Bw*BSH1 was preincubated with 0, 0.1, 1, 10, 100, and 1000 µM BSH-IN-1 (DMSO stock solution) or 0, 0.1, 1, 10, 100, and 1000 µM CHA (MeOH stock solution) for 30 min in PBS (pH 7.4). TCA (1 mM) was added to each solution, and the resulting mixtures were incubated at 37 °C. At 15 min timepoint, 10 μL of each reaction mixture was subsequently quenched by adding 90 μL of ACN followed by centrifugation (16,000 × *g* for 15 min) to remove the precipitated protein. The reaction was monitored by UPLC-MS/MS as described in **UPLC-MS/MS analytical methods**. Enzyme activity was reported as the percentage of the initial activity.

### Activity-guided purification of *Bw*BSHs from *B. wadsworthia* culture medium

*Preparation of the supernatant*: Four 5 mL cultures of ABB supplemented with 5 mM taurine in a Hungate tubes were inoculated from a *B. wadsworthia* 3_1_6 glycerol stock. The starter cultures were grown at 37 °C to saturation. Next, four 500 mL volumes of ABB supplemented with TCA were each inoculated with a 5 mL starter culture, sealed, and incubated at 37 °C. After 24 h of growth, the cultures were centrifuged at 5,000 × *g* for 10 min at 4 °C. The cell pellets were discarded, and the resulting supernatants was combined and filtered through a 0.2 μm membrane prior to purification.

*Ammonium sulfate precipitation and desalting*: The filtered supernatant (1.9 L) was fractionated using the slow addition of solid ammonium sulfate to reach 40% saturation while stirring gently at 4 °C. After 60 min of stirring at 4 °C, the precipitated proteins were pelleted by centrifugation (6,000 × *g*, 20 min) and dissolved in 50 mL of HEPES buffer (20 mM HEPES, 100 mM NaCl, pH 8.0). The remaining supernatant was sequentially brought to 60% and then 80% ammonium sulfate saturation, with the resulting pellets collected and dissolved in the same manner. Each dissolved protein fraction was dialyzed to remove residual ammonium sulfate and concentrated using Amicon Ultra centrifugal filters (50 kDa MW). The concentrates were assayed for BSH activity. The highest activity was observed in the fraction precipitated at 60% saturation.

*Size-Exclusion Chromatography (SEC)*: The active fraction from the 60% ammonium sulfate precipitation was concentrated to a volume of 3 mL and loaded onto a HiLoad 16/600 Superdex 200 pg column pre-equilibrated with HEPES Buffer (20 mM HEPES, 100 mM NaCl, pH 8.0). The FPLC trace at 280 nm appeared as a single broad peak, likely due to background absorbance caused by *B. wadsworthia* sulfide production. Protein purity was assessed by SDS-PAGE using Novex 4–20% Tris-Glycine Plus WedgeWell gels. The gel band corresponding to the fraction with the highest BSH activity was excised and subjected to proteomic analysis. The initial BSH activity of native secretory *Bw*BSHs from *B. wadsworthia* 3_1_6 was evaluated by comparison with recombinant *Bw*BSH1 (WP_005025869) at equivalent protein concentrations.

#### Structure prediction and docking

We utilized a local installation of AlphaFold3 (v3.0.1) to predict the structure of *Bw*BSH1. The enzyme was co-folded with two Zn^2+^ ions and TCA as the substrate, which yielded the structure of the holoenzyme. The AlphaFold3 input required a SMILES string for TCA, which was obtained after building the ligand in Avogadro 2.0^96^ with a charge of −1 on the sulfonate group. Out of six predicted protein models from AlphaFold3, we selected the one where the Zn^2+^ ions were coordinated (i.e., as judged by a heuristic metal-ligand bond distance cutoff of 2.3 Å) by the histidine (H110, H112, H441, H474), aspartate (D583), and carboxylated lysine (KCX373) residues. We also targeted a Zn^2+^-Zn^2+^ distance of 3.95 Å, which is the distance in *Caulobacter crescentus* dipeptidase (PDB: 3MTW). The TCA ligand was then removed from the AlphaFold3-predicted receptor.

The best AlphaFold3-predicted receptor was protonated using Protoss^97^, as implemented in the *qp.protonate* module of the QuantumPDB package^98^. The protonation states of the residues in the core active site were reviewed and manually reassigned, where necessary, such that histidine residues (H110, H112, H441, H474, H196) were neutral, and the Zn^2+^-coordinating aspartate residue had a charge of −1 (**Supplementary Table 16**). The SMILES string of TCA was then used to protonate the ligand using the Molscrub python package^99^. PDBQT files for both the receptor and the ligand were generated using the *prepare_receptor* and *mk_prepare_ligand.py* utilities from the ADFR Suite v1.0^100,101^ and AutoDock Vina^57,58^ respectively.

For the docking experiments, AutoDock Vina 1.2.7 was employed. The docking grid box was centered on the following residues: α-Zn^2+^, β-Zn^2+^, H110, H112, H196, KCX373, H441, H474, and D583. The geometric center (center_x = −3.317 Å, center_y = −1.311 Å, center_z = 11.040 Å) was determined through the *centroid.py* plugin for PyMOL3. The docking box had dimensions of 25 x 25 x 25 Å^3^. The exhaustiveness parameter was set to 32, the energy range was set to 10, and the number of modes was set to 20.

The best docking pose was considered to be the one that had the lowest DFT energy. It also satisfied the following criteria: i) the sulfonate group of the substrate is deep into the active site and ii) the Zn^2+^-carbonyl oxygen distance was not below 2 Å, chosen based on the sum of covalent radii of the two atoms.

To obtain an initial guess of the orientation of the hydroxide within the active site, we performed a separate, hydrated docking experiment with AutoDock Vina 1.2.7 using the AutoDock4 forcefield^102^. Affinity maps for the receptor were generated with the *mk_prepare_receptor.py* utility, where the box parameters were the same as the previous docking experiments. The ligand was decorated with an ensemble of water molecules using the mk_prepare_ligand.py script. Affinity maps for water and all other water types were generated using the output grid parameter file from *mk_prepare_receptor.py*. For this docking experiment, the exhaustiveness parameter was set to 32, the energy range was set to 10, the number of modes was set to 20, and the AutoDock4 scoring function was employed.

#### Quantum mechanical (QM) cluster model preparation and calculations

To assess the best binding pose from the docking experiments and optimize the geometry of the active site, we constructed large QM cluster models^103^ that were comprised of the TCA ligand, Zn^2+^ ions, and all protein residues surrounding them within the first two coordination spheres, constructed with *qp.cluster* module of the QuantumPDB package using a Voronoi tessellation routine. The cutoff distance for merging individual cluster centers was set at 4.5 Å. Any truncated peptide bonds in the cluster were capped with hydrogen atoms. This process resulted in 750-atom-large clusters, with a net charge of +1 (**Supplementary Table 17**). For each cluster built from a docking pose, we performed a density functional theory (DFT) single-point energy calculations using the developer version 1.96H of the GPU-accelerated quantum chemistry package TeraChem^104–107^. These closed shell singlet calculations were carried out with the ωB97X-D3(BJ) functional^108,109^ and a basis set corresponding to the LANL2DZ effective core potential for zinc and 6-31G* applied to all other atoms. A modest double zeta basis set was chosen due to the 750-atom size of the QM cluster. A level shift of 0.25 Ha was applied for virtual orbitals. Solvation effects were included using the C-PCM model^110,111^ as implemented in TeraChem, with a dielectric constant of 10 to mimic the protein environment.

The QM cluster model with the best pose for TCA was subjected to geometry optimization using ωB97X-D3(BJ)/6-31G* as in the single-point calculations. The geomeTRIC optimizer^112^ was used, as implemented in TeraChem. All backbone atoms of the active pocket residues (i.e., alpha carbon, carboxyl carbon and oxygen, as well as nitrogen) were fixed in place in order to mimic the enzyme scaffold around the active site and prevent the unphysical movements of the residues. The geometry optimization was converged using thresholds of 1.0 × 10^-6^ Ha for the energy, 3.0 × 10^-4^ and 4.5 × 10^-4^ Ha·bohr^-1^ for the RMS and maximum gradients, and 1.2 × 10^-3^ and 1.8 × 10^-3^ bohr for the RMS and maximum atomic displacements, respectively. A Zenodo repository containing the AlphaFold3-generated receptor mode, the docking poses, the QM cluster model, and the optimized geometry of the active site is available at Zenodo (record 19406152).

#### Sequence Similarity Networks (SSNs) analysis

To construct an SSN of amidohydrolases, including *Bw*BSHs we first downloaded 110,952 amidohydrolase family protein sequences (Pfam: PF07969)^32^ from the UniProtKB database^113^ (accessed July 13, 2025). To supplement this list, we queried the NCBI non-redundant (nr) database with the *Bw*BSH1 protein sequence (WP_005025869) via BLASTP, identifying 5,000 *Bw*BSH homologs and related amidohydrolases (*E* ≤ 8 × 10^-49^). Additionally, we used Protrek^114^ to search the Open MetaGenomic (OMG) corpus^115^, yielding another 5,000 *Bw*BSH homologs and related amidohydrolases (*E* ≤ 7 × 10^-13^). These three sequence sets were combined and deduplicated using SeqKit v2.10.1^116^, yielding an initial dataset of 112,014 unique sequences. We then scanned this dataset against the Pfam hidden Markov model (HMM) library using HMMER 3.1b2^117^, retaining 101,406 non-fragment Amidohydro_3 family sequences for downstream analysis. To reduce redundancy, sequences were grouped at 65% aa identity using MMseqs2 v18.8cc5c^118^. For the SSN analysis, we selected only the representative sequences from each group alongside functionally annotated marker sequences, retaining the sequence-mapping data for reference. Ultimately, this curated list of 21,230 sequences was submitted to the Enzyme Function Initiative’s Enzyme Similarity Tool (EFI-EST)^119^. To determine the optimal alignment score, networks were generated across a range of thresholds. The final cutoff was selected to ensure the proper clustering of identified metalloBSHs and characterized amidohydrolases within this Pfam family. Cytoscape v3.10.4 was used to visualize and refine the network layout^120^. Documentation of the SSNs for the PF07969 amidohydrolase family is available in a Zenodo repository (record 19406152). The repository contains the protein sequences, representative node sequences and their mappings, SSNs constructed at different alignment scores, and the metalloBSH sequences.

### Transcriptional analysis of *B. wadsworthia* ATCC 49260

To analyze the in vivo transcriptome of *B. wadsworthia* ATCC 49260, metatranscriptomic data were acquired from the NCBI database (PRJNA445875)^28^. The dataset consisted of total RNA extracted from the caeca of altered Schaedler flora (ASF) mice that were colonized with *B. wadsworthia* (N = 4) or co-colonized with *Lactobacillus rhamnosus* and *B. wadsworthia* (N = 5). To detect *B. wadsworthia*-specific transcripts, reads were mapped against its reference proteome. For clustered protein sequences, the representative sequence was defined as the sequence possessing the highest total alignment score within its cluster. To prevent non-specific mapping, the reference database was supplemented with proteomes from the eight ASF strains (ASF 356, 360, 361, 457, 492, 500, 502, and 519). Alignment was performed using DIAMOND BLASTX (v2.1.13)^121^ with an e-value cutoff of 0.0001 and the ’mid-sensitive’ profile. Only reads demonstrating > 90% aa identity to a target sequence were counted. Gene expression levels were quantified as transcripts per million (TPM) and log-transformed using log_10_(TPM+1). Data visualization was performed using GraphPad Prism v10.2.0.

#### MetalloBSH prevalence and metabolomic correlations

*<u>Data Acquisition:</u>* Matched metagenomic and metabolomic datasets were acquired from three distinct cohorts: *Human Intestinal Cohort*^29^ – Metagenomic data from intestinal samples (collected via pH-responsive ingestible devices) and stool samples were downloaded from the NCBI Sequence Read Archive (PRJNA822660). The corresponding metabolomic dataset of amino acid-conjugated bile acids from intestinal and stool samples were retrieved from the bile_salts_wAAconjugates.csv file, available on Zenodo (record 7683655) or via the publication’s GitHub repository (commit 5674501).

*Inflammatory Bowel Disease (IBD) Cohort*^89^ – Stool metagenomic and metabolomic data from the HMP2 Multi’omics Database were obtained from NCBI Sequence Read Archive (PRJNA398089) and the Metabolomics Workbench (PR000639), respectively.

*Colorectal Cancer (CRC) Cohort*^88^ – Stool metagenomic data were downloaded from DDBJ (DRA006684 and DRA008156), and paired metabolomic data were extracted from Supplementary Table S13 of the original manuscript.

*<u>Metagenomic Preprocessing and Gene Quantification:</u>* Host-associated reads, low-quality sequences, and adapters were filtered from the metagenomic data using KneadData (v0.12.3) with default settings. To detect genes encoding metalloBSHs, we curated a reference protein database representing the metalloBSH cluster from our SSN analysis of the amidohydrolase family (PF07969) (**Supplementary Table 9b**). The sequence with the highest sum of alignment scores was selected as the representative reference. To minimize non-specific read mapping, all sequences from the SSN that fell outside the metalloBSH cluster were also included in the database as decoy sequences.

Quality-filtered reads were aligned against this database using a DIAMOND BLASTX search (v2.1.13; e-value cutoff: 0.0001, sensitivity profile: mid-sensitive). Reads exhibiting > 90% amino acid identity to a target sequence were counted. To maximize quantification specificity, multi-mapping reads were exclusively assigned to the protein with the highest identity score. Hit counts were first normalized to Reads Per Kilobase Million (RPKM) by adjusting for total sample read depth and protein length. For accurate cross-sample statistical comparisons, these RPKM values were further normalized by the Average Genome Size (AGS) of each metagenomic sample, estimated using MicrobeCensus ^122^. A metalloBSH-encoding gene was considered “present” in a sample if its AGS-normalized abundance exceeded a threshold of 10^-4^. Differences in metalloBSH prevalence between biological groups were assessed using a chi-square test.

*<u>MGX and MBX Correlation and Visualization:</u>* To evaluate the relationships between *metallobsh* gene abundances and corresponding intestinal or stool metabolite concentrations, only paired metagenomic and metabolomic data were analyzed. Statistical correlations between gene abundance and metabolite levels were assessed on untransformed data using Spearman’s rank correlation coefficient (ρ). To account for multiple comparisons, *P* values were adjusted using the Benjamini–Hochberg false discovery rate (FDR) correction. Statistical analyses were performed using the scipy.stats (v1.16.1) Python module.^123^ For visualization, gene abundances and metabolite concentrations were log_2_- and log_10_-transformed, respectively. To accommodate zero values, a pseudo-count equal to half of the minimum non-zero value in each respective dataset was added prior to transformation.^124^ Scatter plots illustrating these multi-omic associations were generated using the matplotlib (v3.10.7) and seaborn (v0.13.2) libraries,^125,126^ with the resulting correlation coefficients and corrected *P* values annotated directly on the plots.

## Data Availability

Previously published crystal structures are available in the Protein Data Bank (https://www.rcsb.org/) under accession codes 3MTW and 3ICJ. Protein expression vectors are available in Addgene. All other data supporting the findings of this study are available within the manuscript or Supplementary Information. Source data are provided with this paper.

## Code Availability

Data related to *Bw*BSH1 structure prediction and docking (including the AlphaFold3-generated receptor model, docking poses, QM cluster model, and optimized active site geometry), sequence similarity network for the amidohydrolase family PF07969, along with custom code used for bioinformatics analysis and statistical calculations, are available on Zenodo at doi:10.5281/zenodo.19406152.

## Acknowledgments

We thank all members of the Balskus group for insightful discussions. We thank the Taplin Mass Spectrometry Facility at Harvard Medical School for proteomics services. This work was supported by Harvard University and the National Institute of General Medical Sciences (NIGMS) of the National Institutes of Health (NIH) under award number R35GM152027 (for P.P.P., A.T.P.N., and H.J.K.). Individual support was provided by an NIH/NIGMS fellowship (F32GM151795 to Z.C.), an NSERC Postgraduate Scholarship–Doctoral award (to S.M.I.), and an XtalPi AI for Science Fellowship (to A.T.P.N.). E.P.B is a Howard Hughes Medical Institute (HHMI) Investigator. The bioinformatic analyses in this paper were performed on the FASRC Cannon cluster, supported by the FAS Division of Science Research Computing Group at Harvard University. The authors acknowledge the MIT Office of Research Computing and Data for providing high performance computing resources that have contributed to the research results reported within this paper. The authors acknowledge the MIT SuperCloud and Lincoln Laboratory Supercomputing Center for providing HPC resources that have contributed to the research results reported within this paper. Support for the ICP-MS instrument was provided by a core center grant P30-ES002109 from the National Institute of Environmental Health Sciences of the National Institutes of Health.

## Contributions

E.P.B. conceived the project. S.M.I. initiated the study and performed the bile salt hydrolysis assays with *B. wadsworthia* strains. Z.C. and C.J.M. performed the activity-guided native protein purification and identified *Bw*BSH1. Z.C. conducted all other biochemical experiments, identified additional metalloBSHs and short amidohydrolases with BSH activity. Z.C. synthesized cholylhydroxamic acid based on suggestions from R.Y. and M.A.A. P.P.P. and A.T.P.N. performed the molecular docking experiments and quantum mechanical modeling under the guidance of H.J.K. Z.C. and H.E.A. built the protein sequence similarity network for the amidohydrolase family PF07969. Z.C. conducted the bioinformatic analysis of *metallobshs*, with the methodology verified by H.E.A. All authors provided critical feedback on the experiments. Z.C. and E.P.B. wrote the manuscript, which was reviewed and edited by all authors.

---

**Extended Data Fig. 1.**
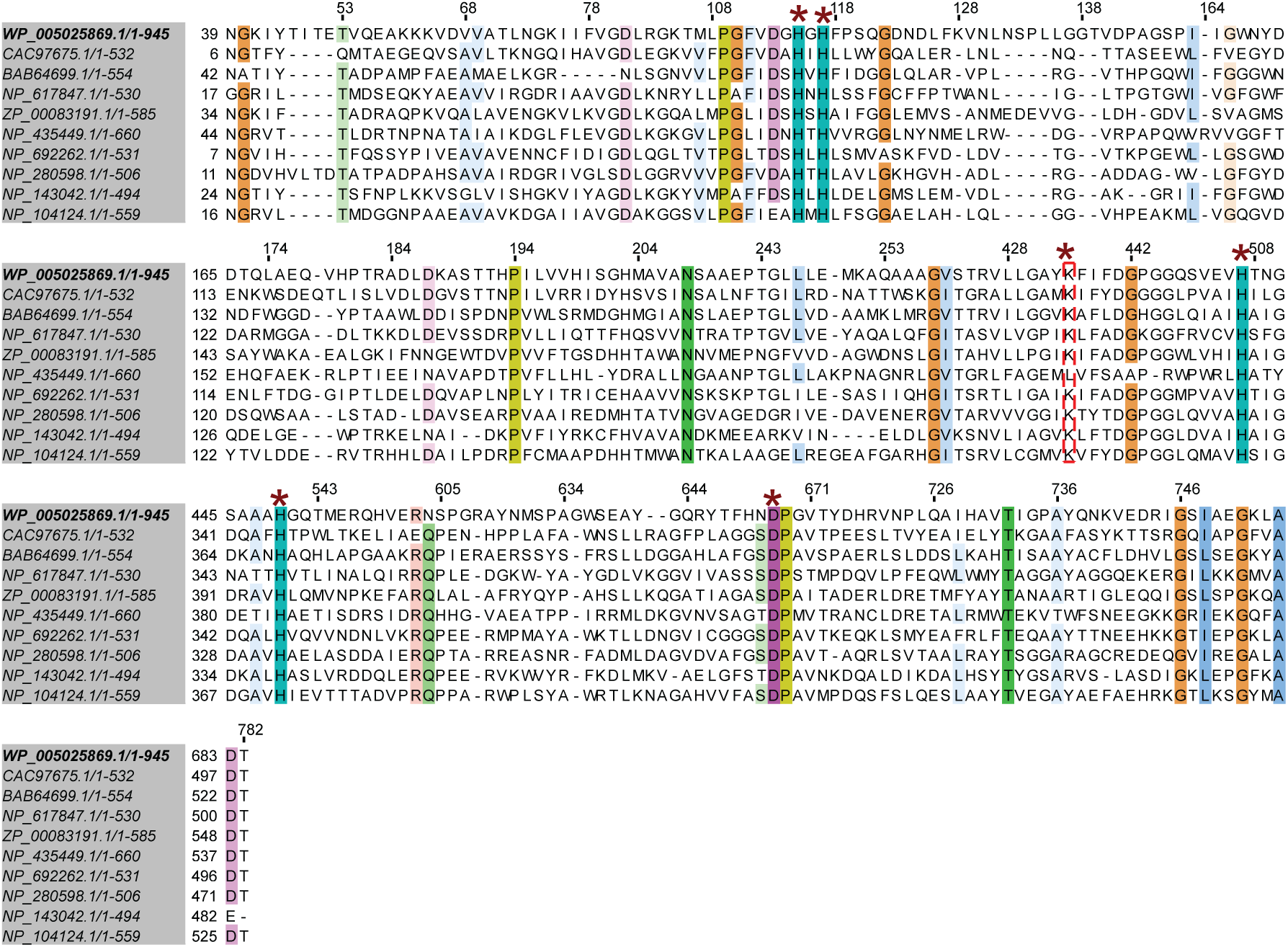
: Multiple sequence alignment of *Bw*BSH1 and YtcJ (Accession: COG1574), YtcJ_like metal dependent amidohydrolases (Accession: cd01300) and amidohydrolase family members (Accession: Pfam07969) identified from a conserved protein domain search. The six conserved residues that are predicted to be involved in metal binding based on similarity to sequences of other amidohydrolase family members are highlighted with *.

**Extended Data Fig. 2 :**
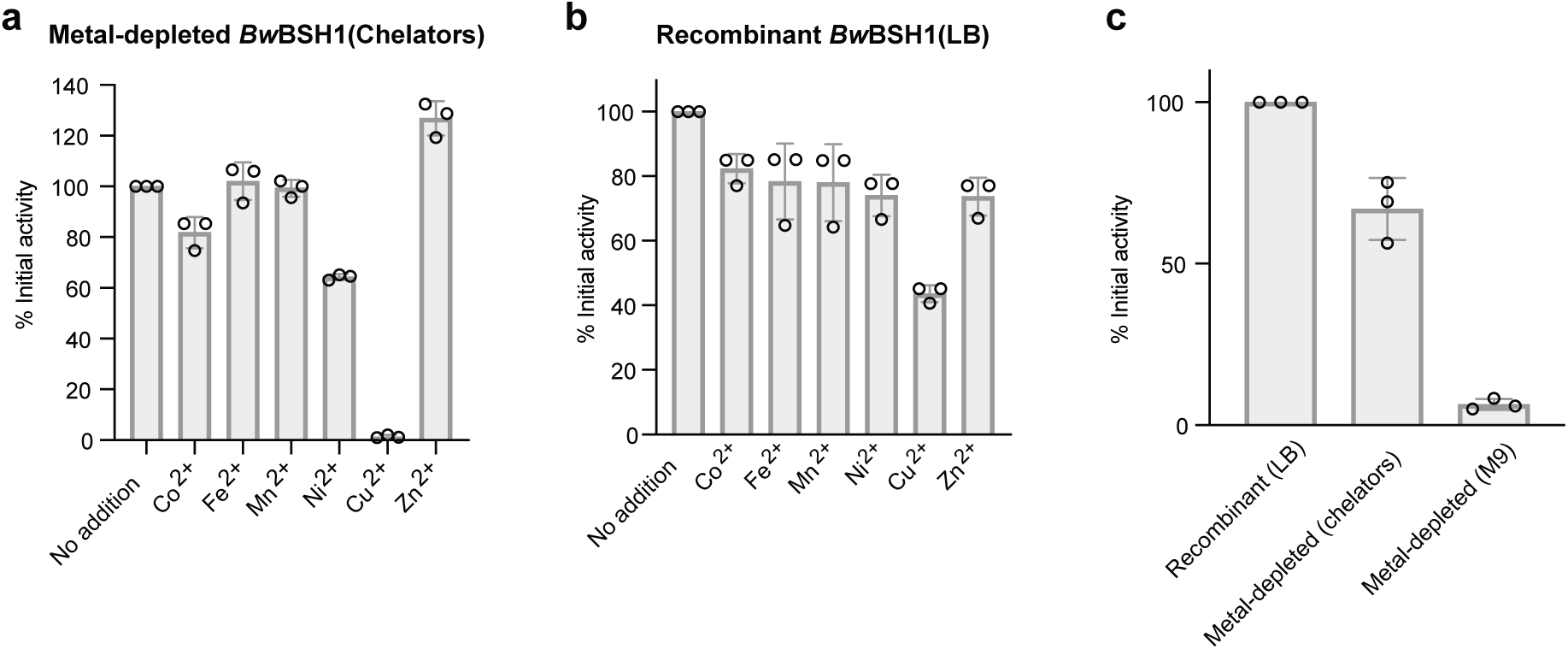
Impact of metal supplementation on *Bw*BSH1 initial activity. a, Initial activity of chelator-treated *Bw*BSH1 in the presence of various metals. Cu²⁺ inhibits activity, while Zn²⁺ enhances it. b, Initial activity of LB-purified *Bw*BSH1 with metal supplementation, highlighting inhibition by Cu²⁺. c, Comparison of basal initial activity across *Bw*BSH1 preparations. The enzyme expressed in metal-depleted M9 medium exhibits lower activity than both chelator-treated and LB-purified recombinant *Bw*BSH1. For panels a, b and c, bars represent the mean of three biological replicates, and error bars indicate the standard deviation.

**Extended Data Fig. 3:**
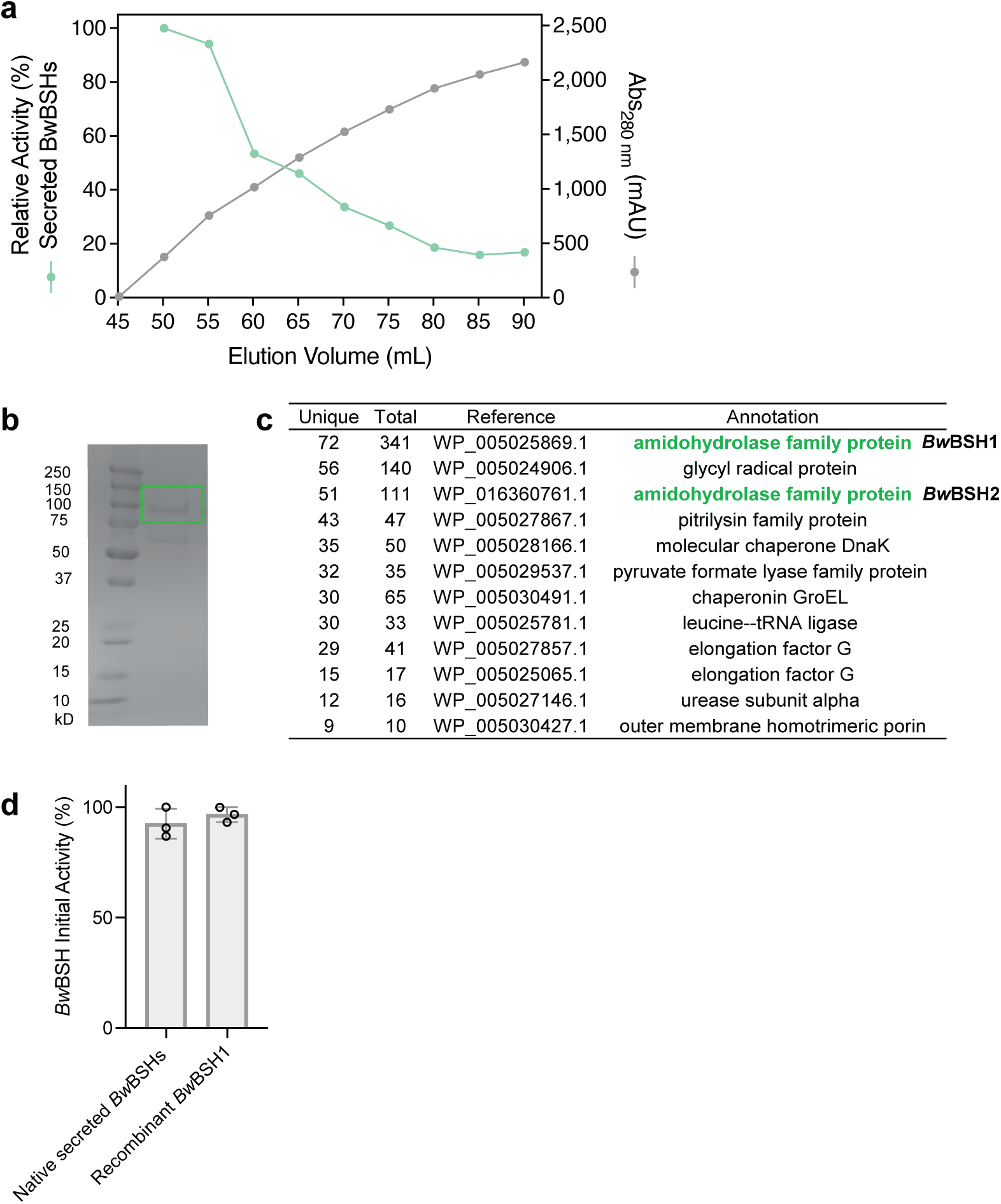
Activity-guided purification of native secreted *Bw*BSHs from *B. wadsworthia* 3_1_6 culture medium. a, Relative TCA hydrolysis activity and UV absorbance (280 nm) of fractions following ammonium sulfate precipitation. b, SDS-PAGE of pooled active fractions. The circled band was excised for proteomic analysis. c, Proteomic results confirming the presence of both *Bw*BSH1 and *Bw*BSH2 in the excised band. A complete list of detected proteins is provided in Supplementary Table 4. d, Activity assays demonstrating comparable BSH activity between natively purified *Bw*BSHs and recombinant *Bw*BSH1. Bars represent the mean of three biological replicates, and error bars indicate the standard deviation.

**Extended Data Fig. 4:**
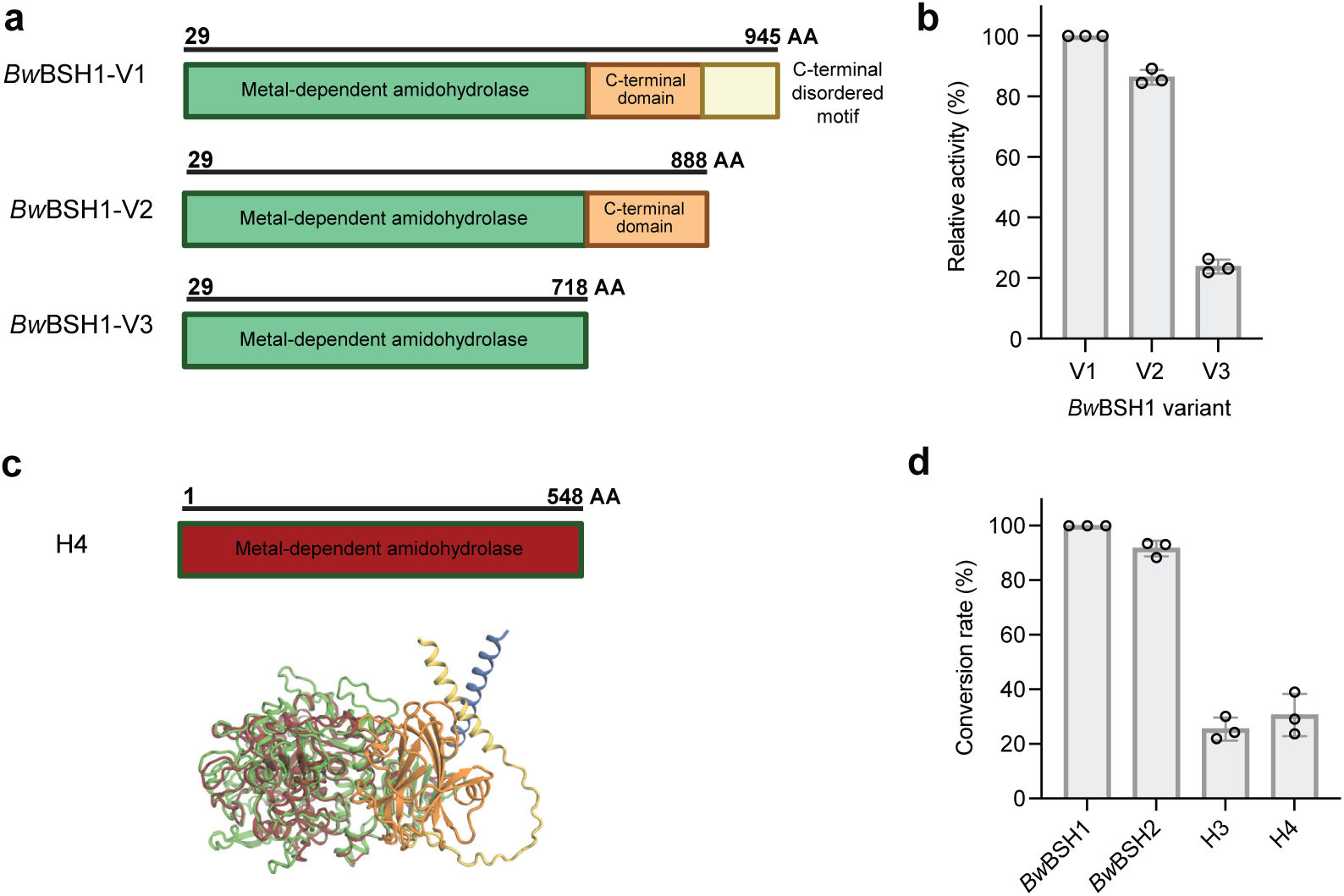
Impact of the additional C-terminal domain on *Bw*BSH activity. a, Domain architecture of *Bw*BSH variants. b, Initial activity of *Bw*BSH1 variants containing different C-terminal domain constructs. c, Domain organization of the short *Bw*BSH1-like amidohydrolase H4 (WP_016360969). Structural alignment of the AlphaFold3-predicted structures of H4 and *Bw*BSH1 (WP_005025869) highlights the absence of the C-terminal domain in short amidohydrolases. d, Total conversion rate of TCA hydrolysis catalyzed by *Bw*BSH1, *Bw*BSH2, H3 (WP_005026994) or H4. Conversion rate (%) is expressed as the percentage of taurine product generated by each enzyme compared to wild-type *Bw*BSH1 following an overnight incubation. For panels b and d, bars represent the mean of three biological replicates, and error bars indicate the standard deviation.

**Extended Data Fig. 5:**
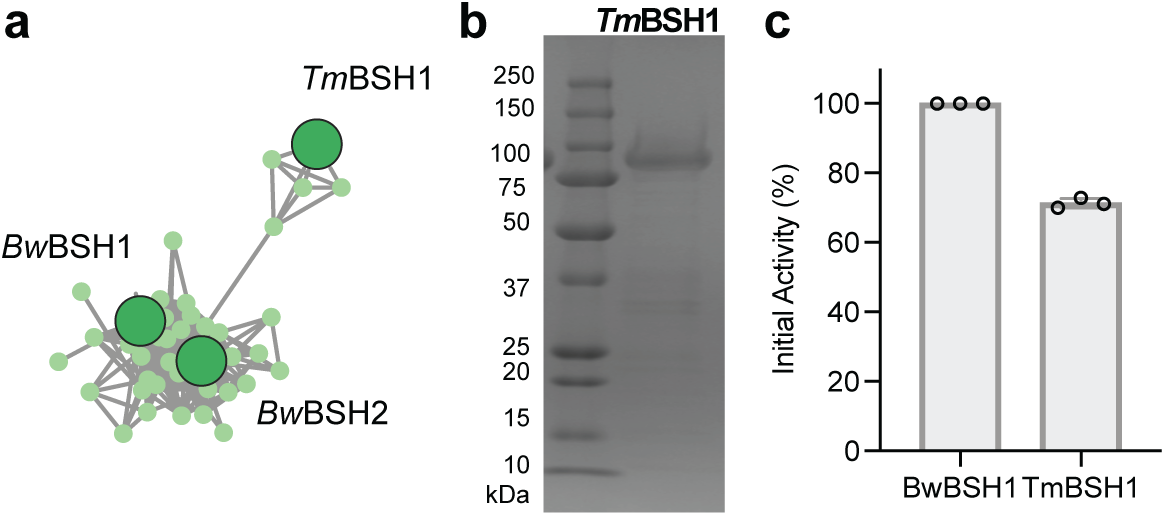
The taurine-respiring mouse gut bacterium *Taurinivorans muris* lacks canonical BSHs but encodes metalloBSHs. a, *Tm*BSH1 groups within a subcluster of metalloBSHs and shares 33.9% aa sequence identity with *Bw*BSH1. b, SDS-PAGE analysis of *Tm*BSH1 (WP_334315222.1, truncated variant; Supplementary Table 9) purified from recombinant *Escherichia coli* BL21(DE3) cultures. The theoretical molecular weight of the N-His_6_-tagged *Tm*BSH1 truncated variant as calculated by ExPASy ProtParam, is 96.5 kDa. c, Activity assays demonstrating comparable BSH activity between recombinant *Bw*BSH1 and *Tm*BSH1. Bars represent the mean of three biological replicates, and error bars indicate the standard deviation.

**Extended Data Fig. 6:**
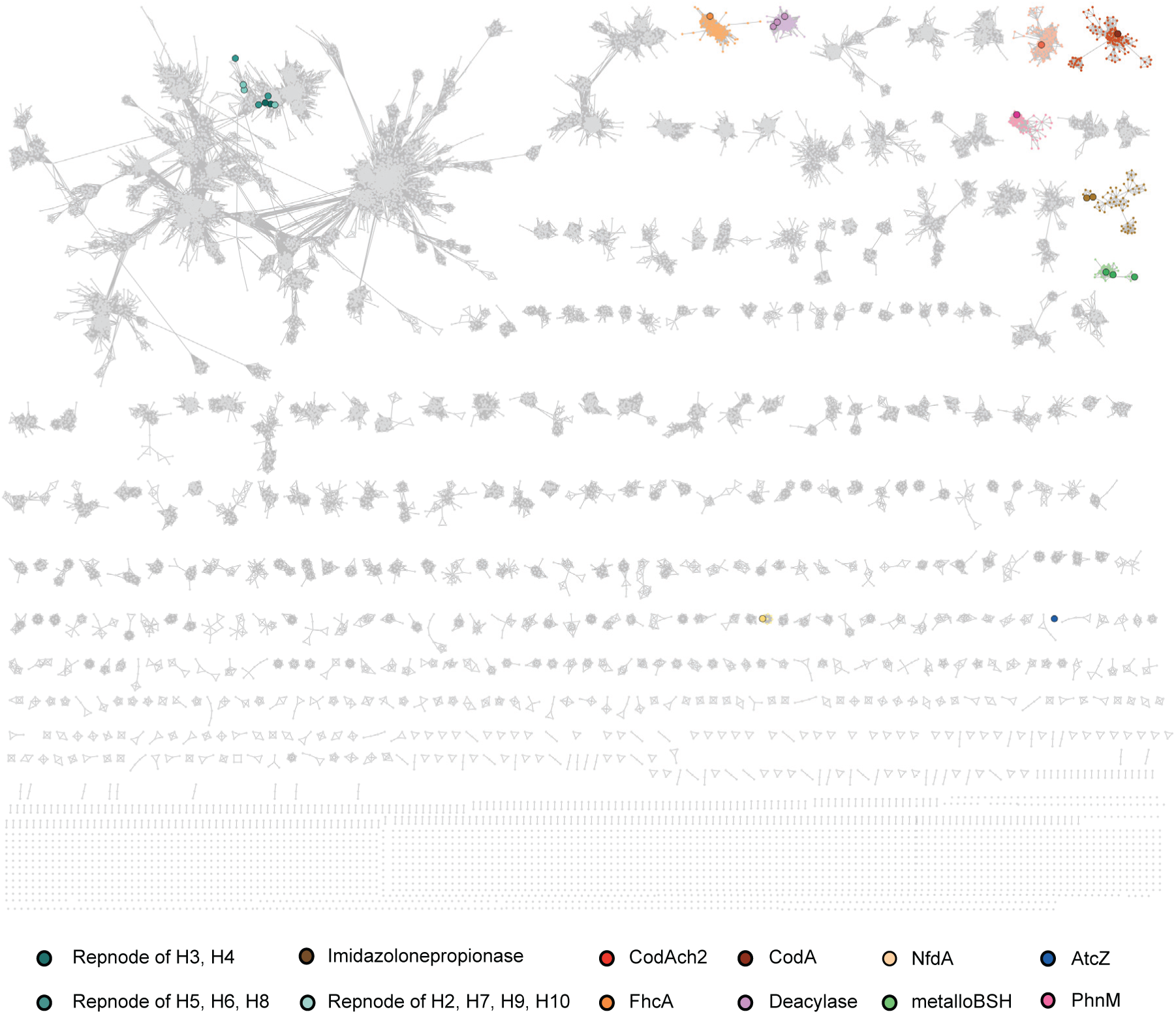
Sequence similarity network (SSN) of amidohydrolase family PF07969. Nodes representing uncharacterized sequences are shown in light gray. Nodes representing characterized sequences are distinctly colored, and experimentally validated enzymes are represented by enlarged nodes. The shorter amidohydrolases from *B. wadsworthia* that lack C-terminal domain and possess weak BSH activity (H3, H4, H5, H6 and H8) or no detectable BSH activity (H2, H7, H9 and H10) segregate into a distinct uncharacterized cluster.

**Extended Data Fig. 7:**
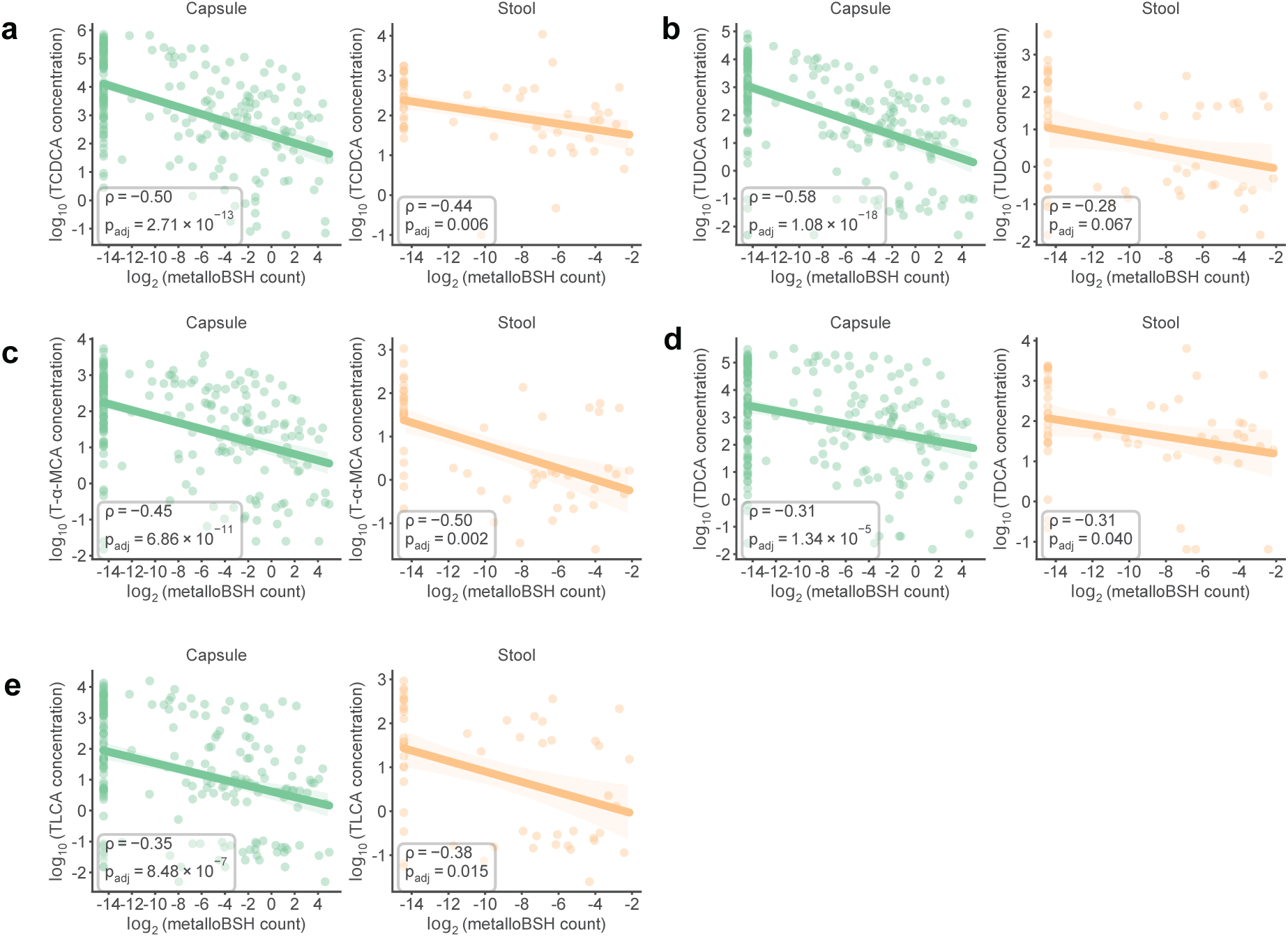
*Metallobsh* gene abundance inversely correlates with taurine-conjugated BA levels. The log_2_(read count) of MetalloBSH-encoding genes in intestinal or stool metagenomes was negatively correlated with levels of all detected taurine-conjugated bile acids in intestinal and stool metabolomes. Correlations are Spearman correlations with Benjamini–Hochberg correction. Data points are individual intestinal (N = 209) or stool samples (N = 56) for which both metagenome sequencing data and metabolomics data were available.

**Extended Data Fig. 8:**
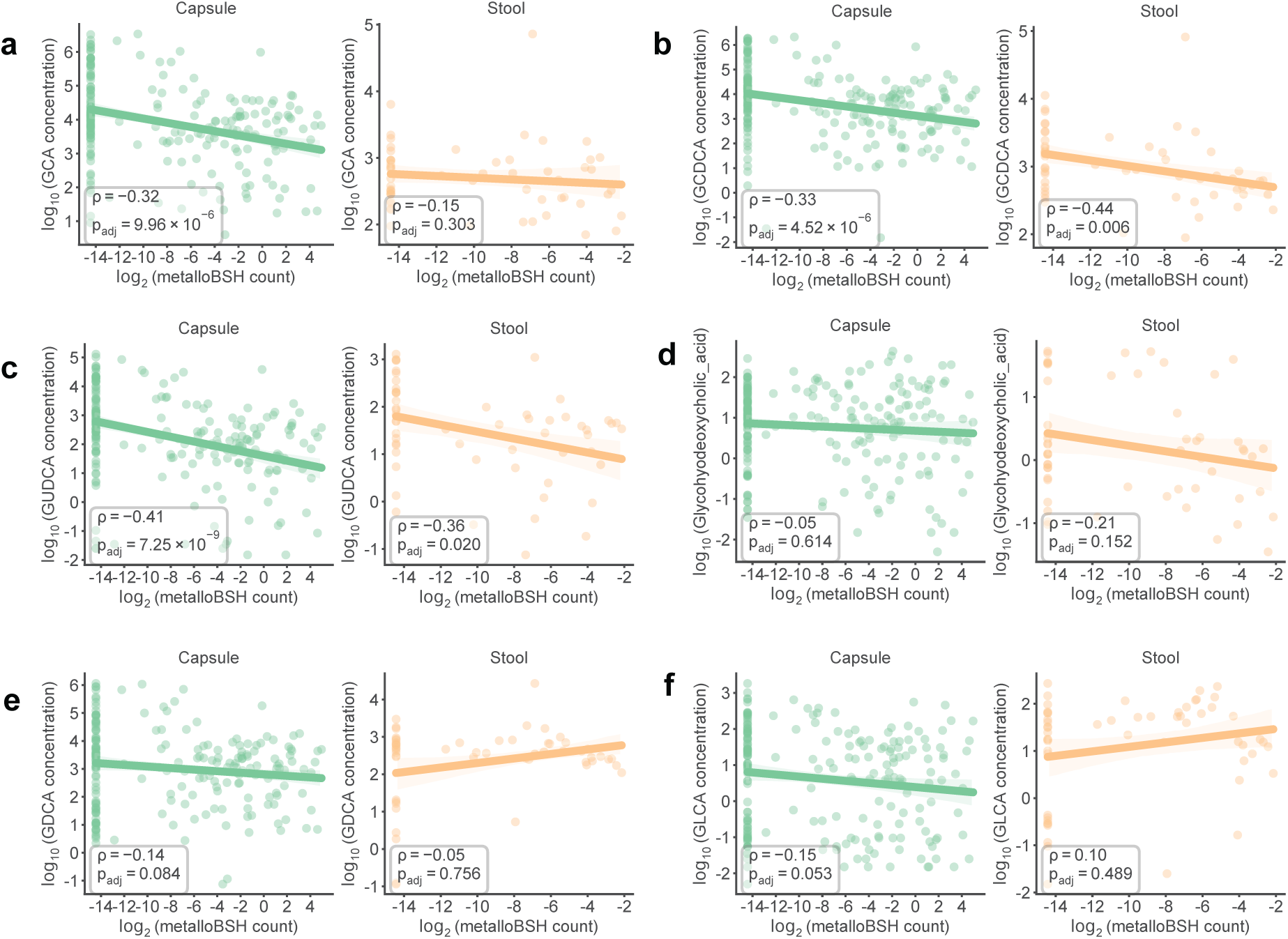
*Metallobsh* gene abundance exhibits divergent correlations with the levels of specific glycine-conjugated BAs in intestinal or stool metagenomes. The log_2_ (read count) of metalloBSH-encoding genes in intestinal or stool metagenomes was divergently correlated with levels of glycine-conjugated bile acids in intestinal and stool metabolomes. Correlations are Spearman correlations with Benjamini–Hochberg correction. Data points are individual intestinal (N = 209) or stool samples (N = 56) for which both metagenome sequencing data and metabolomics data were available.

**Extended Data Fig. 9.**
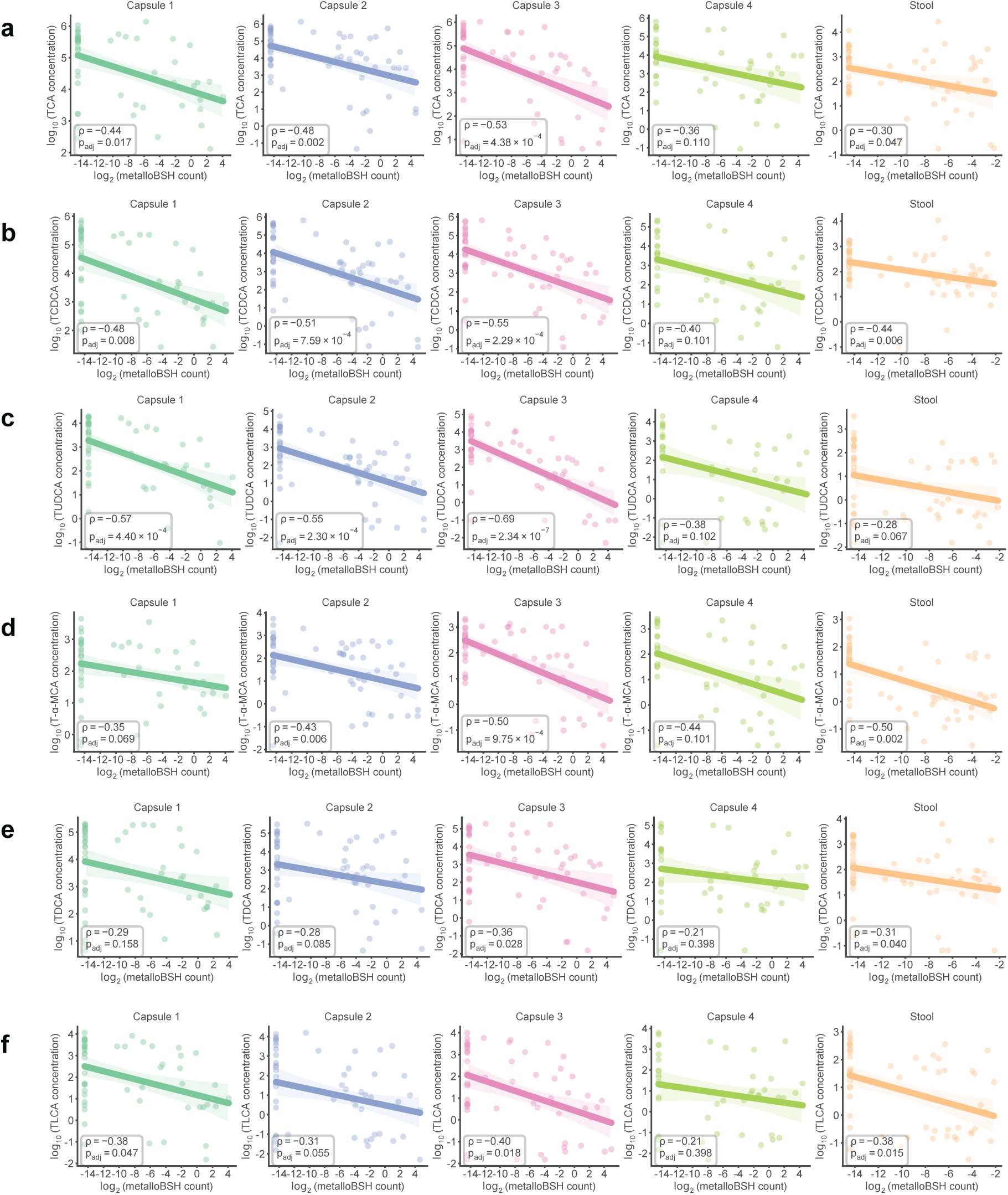
: *Metallobsh* gene abundance inversely correlates with taurine-conjugated BA levels in the small intestine and stool. The log_2_ (read count) of metalloBSH-encoding genes in metagenomes from the pyloric sphincter (Capsule 1), small intestine (Capsule 2 and 3), and stool negatively correlated with nearly all detected taurine-conjugated BAs in paired metabolomes. These inverse correlations are highly significant for TCA (9a), TCDCA (9b), TUDCA (9c), and T-⍺-MCA (9d), with weaker or non-significant correlations observed for TDCA (9e, Capsule 2 *P_adj_* = 0.085) and TLCA (9f, Capsule 2, *P_adj_* = 0.055). Conversely, no significant correlations between metalloBSH abundance and taurine-conjugated BA levels were observed in the ascending colon (Capsule 4). Statistics reflect Spearman correlations with Benjamini–Hochberg correction. Data points represent individual samples with paired metagenomic and metabolomic data (Capsule 1, N = 51; Capsule 2, N= 59; Capsule 3, N = 55; Capsule 4, N = 44; Stool, N =56).

**Extended Data Fig. 10:**
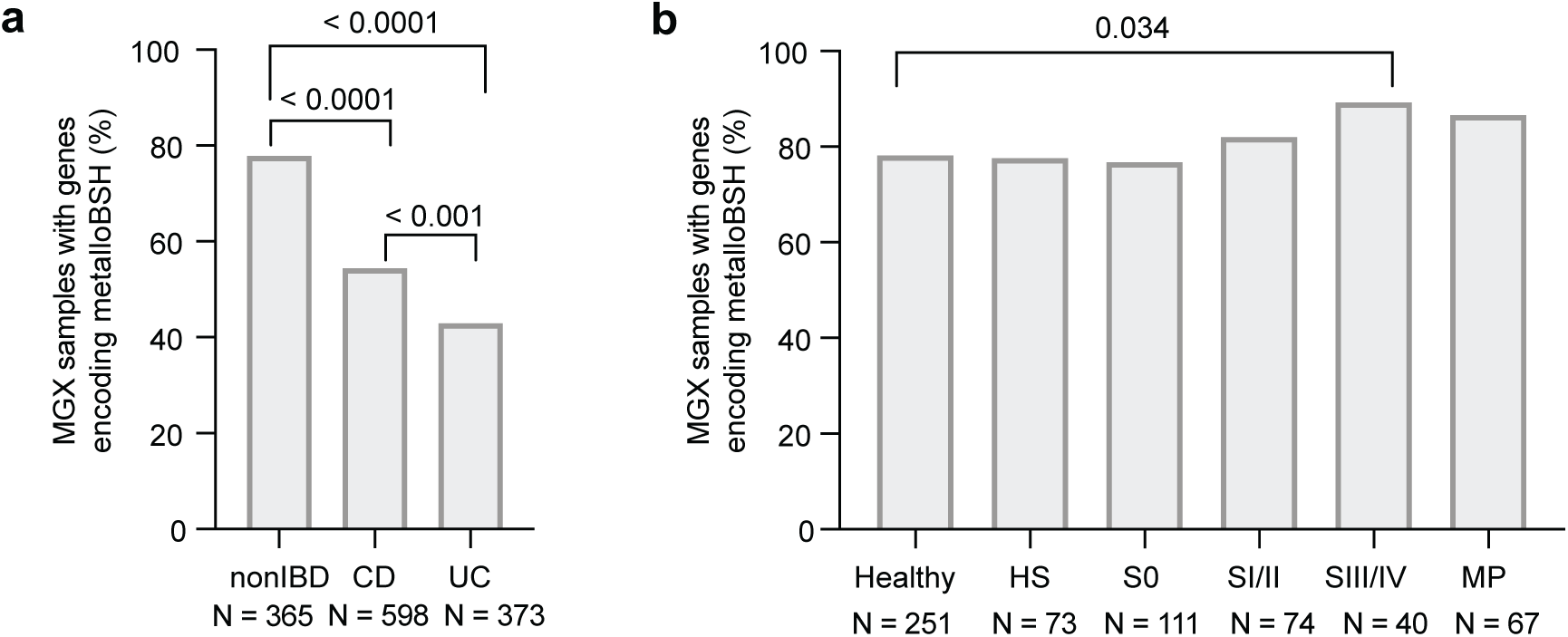
Prevalence of metalloBSHs in the human gut in two disease cohorts. The bar plots show the percentage of MGX samples encoding detectable metalloBSHs across distinct disease groups within (a) the HMP2 cohort and (b) a CRC cohort. The number of metagenomic samples included is provided beneath each corresponding subgroup. A chi-square test of equal proportions was utilized to determine whether the fraction of samples with detectable metalloBSHs differed significantly between groups. *P* values are displayed above the comparisons, unadjusted for multiple comparisons.

## Notes

https://doi.org/10.5281/zenodo.19406152

### Competing Interest Statement

The authors have declared no competing interest.

